# Interleukin-33 coordinates a microglial phagocytic response and limits corticothalamic excitability and seizure susceptibility

**DOI:** 10.1101/2021.08.05.455250

**Authors:** Rafael T. Han, Ilia D. Vainchtein, Johannes C. M. Schlachetzki, Frances S. Cho, Leah C. Dorman, Tessa Johung, Eunji Ahn, Jerika T. Barron, Hiromi Nakao-Inoue, Akshaj Joshi, Ari. B Molofsky, Christopher K. Glass, Jeanne T. Paz, Anna V. Molofsky

## Abstract

Microglia are key remodelers of neuronal synapses during brain development, but the mechanisms that regulate this process and its ultimate impact on neural circuit function are not well defined. We previously identified the IL-1 family cytokine Interleukin-33 (IL-33) as a novel mediator of microglial synapse remodeling. Here we define the phagocytic program induced in microglia in response to IL-33. We find that IL-33 markedly alters the microglial enhancer landscape and exposes AP-1 transcription factor sites that promote target gene expression. We identify the scavenger receptor MARCO and the pattern recognition receptor TLR2 as downstream mediators of IL-33 dependent synapse engulfment. Conditional deletion of IL-33 in the CNS or its receptor on microglia results in increased numbers of excitatory synapses in the corticothalamic circuit and spontaneous epileptiform activity as well as increased seizure susceptibility by early adulthood. These findings define novel mechanisms through which IL-33 coordinates experience-dependent synaptic refinement to restrict hyperexcitability in the developing brain.

## Introduction

Innate immune signaling shapes tissue development and homeostasis, including the remodeling of neuronal synapses in the central nervous system (CNS). Immune dysfunction is also implicated in the pathogenesis of neurodevelopmental disorders including epilepsy^1–3^, autism, and schizophrenia^4^. Microglia are the dominant immune cells in the brain parenchyma and therefore a potential mechanistic link between innate immunity and neurodevelopmental disease. Microglia can both engulf synapses during development and promote new synapse formation^5, 6^. Microglial function is shaped by an exquisite sensitivity to environmental cues that can rapidly alter microglial identity^7–11^. However, how microglia coordinate synapse remodeling in response to environmental cues is not well-defined.

We recently identified the IL-1 family member Interleukin-33 (IL-33) as a novel regulator of microglial synapse remodeling^12, 13^. Microglia are the primary CNS-resident cells that respond to IL-33 via its obligate co-receptor *Il1rl1*, and global deletion of *Il33* or *Il1rl1* during development results in defective microglial engulfment of excitatory synapses. The thalamus is one of the first brain regions to express IL-33^13^ and its expression increases coincident with synapse maturation in this region^14–17^. Increased excitation in the reciprocal connections between thalamus and cortex is one well-described circuit that can drive seizures^18–21^, including a particular type of childhood epilepsy characterized by absence seizures^22^. This raises the question of how IL-33’s function in corticothalamic maturation might impact seizure susceptibility.

In this study, we defined the impact of IL-33 on microglial gene expression, epigenomic landscape, and function. We identified FOS, a component of the AP-1 transcription factor complex, as a regulator of IL-33 dependent target gene expression, and found that the scavenger receptor MARCO and the pattern recognition receptor TLR2 are two downstream regulators of IL-33 dependent phagocytic responses. Loss of CNS-derived IL-33 or its receptor (IL1RL1) on microglia and myeloid cells led to excess excitatory synapses and decreased inhibitory synapses, but did not alter synaptic strength. Mice lacking CNS-derived IL-33 had an increased incidence of spike wave discharges, a characteristic feature of absence seizures, as well as increased susceptibility to convulsive seizures. These data reveal novel mechanisms by which IL-33 promotes microglial phagocytic capacity and defines a functional requirement for IL-33 in refinement of corticothalamic synapses and in restricting seizure susceptibility.

## Results

### IL-33 induces a microglial phagocytic program that includes pattern recognition and scavenging responses

To define how IL-33 impacts microglial function, we first examined its effect on the microglial transcriptome. We performed single-cell sequencing of flow-sorted thalamic CD45+ cells, which were predominately microglia, 4 hours after intracerebroventricular (i.c.v.) administration of 40 ng of recombinant IL-33 or vehicle (**Fig. 1a; Fig S1a**). This dose and time period were chosen to capture the initial cytokine response and did not result in noticeable infiltration of myeloid or lymphoid cells into the CNS (**Fig. S1b**). To determine whether IL-33 responses required direct signaling to the myeloid lineage and to rule out potential off-target effects, we used *Cx3cr1^CreERT2 23^* to conditionally delete the IL-33 receptor *Il1rl1* in the early postnatal period (tamoxifen: P1, P3, P5). We compared vehicle or IL-33 treated controls (*Cx3cr1^CreERT2+/-^)* to IL-33-treated animals lacking myeloid IL-33R (*Cx3cr1^CreERT2+/-^:Il1rl1^fl/fl^)*.

**Figure 1:**
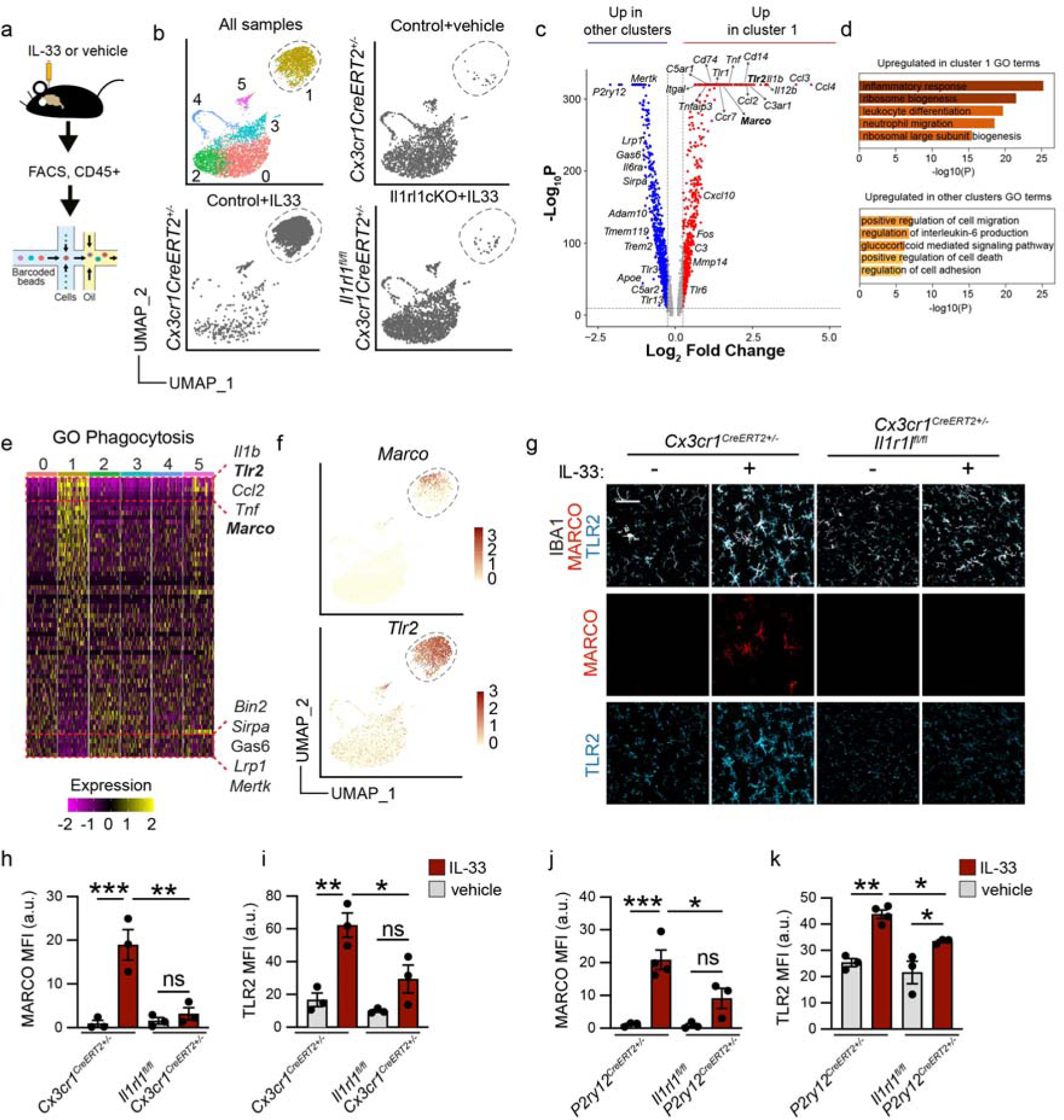
IL-33 induces a microglial phagocytic program that includes pattern recognition and scavenging responses. **a)** Experimental paradigm for single cell RNA-seq from P15 thalamus (vehicle=PBS). **b)** Unsupervised clustering of single cell sequencing data from all three conditions (upper left). Same plot showing only *Cx3cr1^CreERT2^*^+/-^ sample 4 hours after vehicle injection (upper right), showing *Cx3cr1^CreERT2^*^+/-^ sample 4 hours after 40 ng IL-33 injection (lower left), and showing *Cx3cr1^CreERT2^*^+/-^ *Il1rl1^fl/fl^* sample 4 hours after 40 ng of IL-33 injection (lower right) plotted in UMAP space. Each dot represents a cell. **c)** Volcano plot of differentially expressed genes between the IL-33 responsive cluster 1 vs. aggregated cells from all other clusters. Red dots are upregulated in cluster 1 with log_2_ fold change > 0.25, p_Adj_ < 10^-10^, using the MAST test in Seurat. Blue dots are upregulated in all other aggregated clusters vs cluster 1 with log_2_ fold change < 0.25, p_Adj_ < 10^-10^. **d)** Top 5 GO terms upregulated in cluster 1 (upper) and upregulated in all other aggregated clusters (lower). **e)** Heatmap showing expression of phagocytosis related genes (GO:0006909) across clusters, highlighting top 5 upregulated genes (top, yellow) and downregulated genes (bottom, purple) in cluster1 (ordered based on expression in cluster 1, centered normalized expression values). **f)** Feature plots showing *Marco* (upper) and *Tlr2* (lower) expression with cluster 1 highlighted (dotted line). **g)** Representative images of MARCO and TLR2 protein expression in thalamic microglia 18 hours after vehicle or IL-33 injection in *Cx3cr1CreERT2*+/- or *Cx3cr1CreERT2*+/-;*Il1rl1^fl/fl^* mice. Scale bar = 40 µm. **h)** Quantification of MARCO mean fluorescence intensity (MFI) in thalamic microglia 18 hours after vehicle or IL-33 injection in *Cx3cr1CreERT2*+/- or *Cx3cr1CreERT2*+/-;*Il1rl1^fl/fl^* mice. Each dot represents a mouse. Two-way ANOVA followed by Tukey’s post hoc comparison (genotype and treatment). **i)** Quantification of TLR2 mean fluorescence intensity (MFI) in thalamic microglia 18 hours after vehicle or IL-33 injection in *Cx3cr1CreERT2*+/- or *Cx3cr1CreERT2*+/-;*Il1rl1^fl/fl^* mice. Each dot represents a mouse. Two-way ANOVA followed by Tukey’s post hoc comparison (genotype and treatment). **j-k)** Quantification of MARCO **(J)** and TLR2 **(K)** expression in thalamic microglia 18 hours after vehicle or IL-33 injection in *Il1rl1^fl/fl^ or P2ry12creERT2+/-;Il1rl1^fl/fl^* mice. Each dot represents a mouse. Two-way ANOVA followed by Tukey’s post hoc comparison (genotype and treatment). Data represented as mean ± SEM for bar graphs. Mice from P15-P17 were used for g-k. * *p*<0.05, ** *p*<0.01, *** *p*<0.001. ***See also Figure S1 and S2.***

Unsupervised clustering at several resolutions revealed distinct microglial subsets, but only trace levels of macrophages **(Fig. 1b**, quality control in **Fig. S1c-f)**. Most notable was a robust transcriptomic shift in response to IL-33 in 84% of microglia (Cluster 1), indicating that most thalamic microglia are competent to respond to IL-33 signaling. This response was almost completely abrogated after myeloid-specific deletion of the IL-33 receptor (**Fig. 1b**), demonstrating a direct impact of IL-33 on myeloid cells. As expected, we observed an overall activation of immune response pathways. Differential expression analysis of IL-33 responsive cluster 1 vs. all other clusters revealed upregulation of traditional immune activation genes and pathways (*Tnf, Il1b*, GO term: inflammatory response; **Fig. 1c-d**). There was downregulation of homeostatic microglial genes *(P2ry12, Tmem119*). These data indicate that microglia directly respond to IL-33, although we could not rule out an indirect contribution of peripheral, perivascular, or meningeal macrophages to this activation profile.

Phagocytosis is the dominant role of tissue resident macrophages, and we previously demonstrated a role for IL-33 in promoting microglial phagocytosis^12, 13^. To define potential regulatory mechanisms, we correlated IL-33 response genes (cluster 1) with an annotated phagocytosis dataset ((GO: 0006909). This showed that IL-33 induced multiple phagocytic genes while suppressing others (**Fig 1e**). Upregulated genes included scavenger receptors (*Marco, Msr1*) and pattern recognition receptors (*Tlr2*)^24–26^. Some genes linked to phagocytosis of synapses and apoptotic cells were downregulated, including the recognition receptor *Trem2*^27–29^, and TAM receptor tyrosine kinase *Mertk*^30^. Some complement pathway genes were both up and downregulated (up-*C3ar, C5ar*, down-*C5ar2*), consistent with proposed roles of complement in phagocytosis^31, 32^, although the specific roles of these genes in microglia have not been studied. Thus, IL-33 signals directly to myeloid cells and induces a distinct phagocytic gene expression profile which includes many novel functional candidates.

To further examine functional targets of IL-33, we performed immunostaining for two of the top gene candidates in our study: MARCO and TLR2 (**Fig. 1f, Fig. 1g).** The class A scavenger receptor *MARCO*^33^ has been implicated in dendritic cell filopodial morphogenesis and debris clearance^34^ as well as macrophage phagocytosis of bacteria and other particles^35, 36^. We found that MARCO had low baseline expression and was robustly induced in response to IL-33. This induction was completely abrogated after deletion of the IL-33 receptor *Il1rl1* in myeloid cells (*Cx3cr1^CreERT2+/-^:Il1rl1^wt/wt^* vs. littermate *Cx3cr1^creERT2+/-^:Il1rl1^fl/fl^,* tamoxifen at P1,3,5) at the RNA level (**Fig. S1e**) as well as the protein level in both thalamus (**Fig. 1g-h**) and cortex (**Fig. S2g**).

To determine whether induction of MARCO induction requires direct signaling of IL-33 to microglia, as opposed to meningeal, perivascular, or peripheral macrophages, we used a recently created microglial-specific line with Cre recombinase inserted downstream of a self-cleaving peptide in the 3’ end of the microglial-specific gene *P2ry12* (*Il1rl1^+/+^* vs. *P2ry12^creERT2^Il1rl1^fl/fl^* ;tamoxifen at P2, P4, P6^37^). We validated that this strategy was specific to microglia but not perivascular or meningeal macrophages by imaging and flow cytometry (**Fig. S2a-d**). This strategy was somewhat less sensitive, leading to a ∼92% reduction in *Il1rl1* transcript in cortical microglia and an ∼87% reduction in thalamic microglia (**Fig. S2f**). We found that microglial specific deletion of IL-33 receptor significantly reduced MARCO induction after IL-33, both by gene expression (**Fig. S1L**), and at the protein level (**Fig. 1j**, thalamus, **Fig S2i**, cortex). These data indicate that IL-33 signals at least in part directly to microglia to induce expression of MARCO.

Another top functional candidate was TLR2, a pattern recognition receptor that responds to both pathogenic and physiologic signals^38, 39^. TLR2 functions as a heterodimer with TLR1 and TLR6, both of which were also highly expressed in our dataset and induced by IL-33 (**Fig. 1c; Supplemental Table 1**). TLR2 was expressed in all microglia at baseline and was significantly induced by IL-33. Like MARCO, this induction was significantly abrogated after myeloid- or microglia-specific deletion of the IL-33 receptor in both thalamus (**Fig. 1 g, i, k**) and cortex (**Fig. S2h, j**), including at the gene expression level (**Fig. S1e-f**). Taken together, these independent approaches suggest that IL-33 induces target gene expression via direct signaling to microglia.

### MARCO and TLR2 promote IL-33 dependent synapse engulfment and restrict excitatory synapse numbers

We next investigated whether MARCO and TLR2 are causally linked to IL-33’s ability to promote microglial phagocytosis. We previously showed that exogenous IL-33 leads to increased microglial engulfment of synaptic proteins and that it acutely depletes excitatory synapses^13^

We therefore assessed both of these complementary phenotypes after TLR2 or MARCO loss of function. We first quantified microglial engulfment of the presynaptic marker VGLUT1 using high-resolution imaging and 3D reconstruction with the myeloid *Cx3cr1*^GFP^ reporter^40^. This enables more accurate reconstructions of microglial volumes and predominantly labels microglia, as perivascular macrophages are rare in the brain parenchyma (**Fig. S1b, S2a-b**). Injection of IL-33 increased presynaptic protein engulfment, as quantified by the abundance of VGLUT1 within CD68+ microglial phagolysosomes (**Fig. 2a-c**). However, co-injection of IL-33 with either a TLR2 function blocking antibody (**Fig. 2b)**^41, 42^ or a MARCO blocking antibody (**Fig. 2c)**^26, 34^ significantly but partially blunted this response relative to an IgG isotype control. Thus, both MARCO and TLR2 partly mediate IL-33 dependent synapse engulfment.

**Figure 2:**
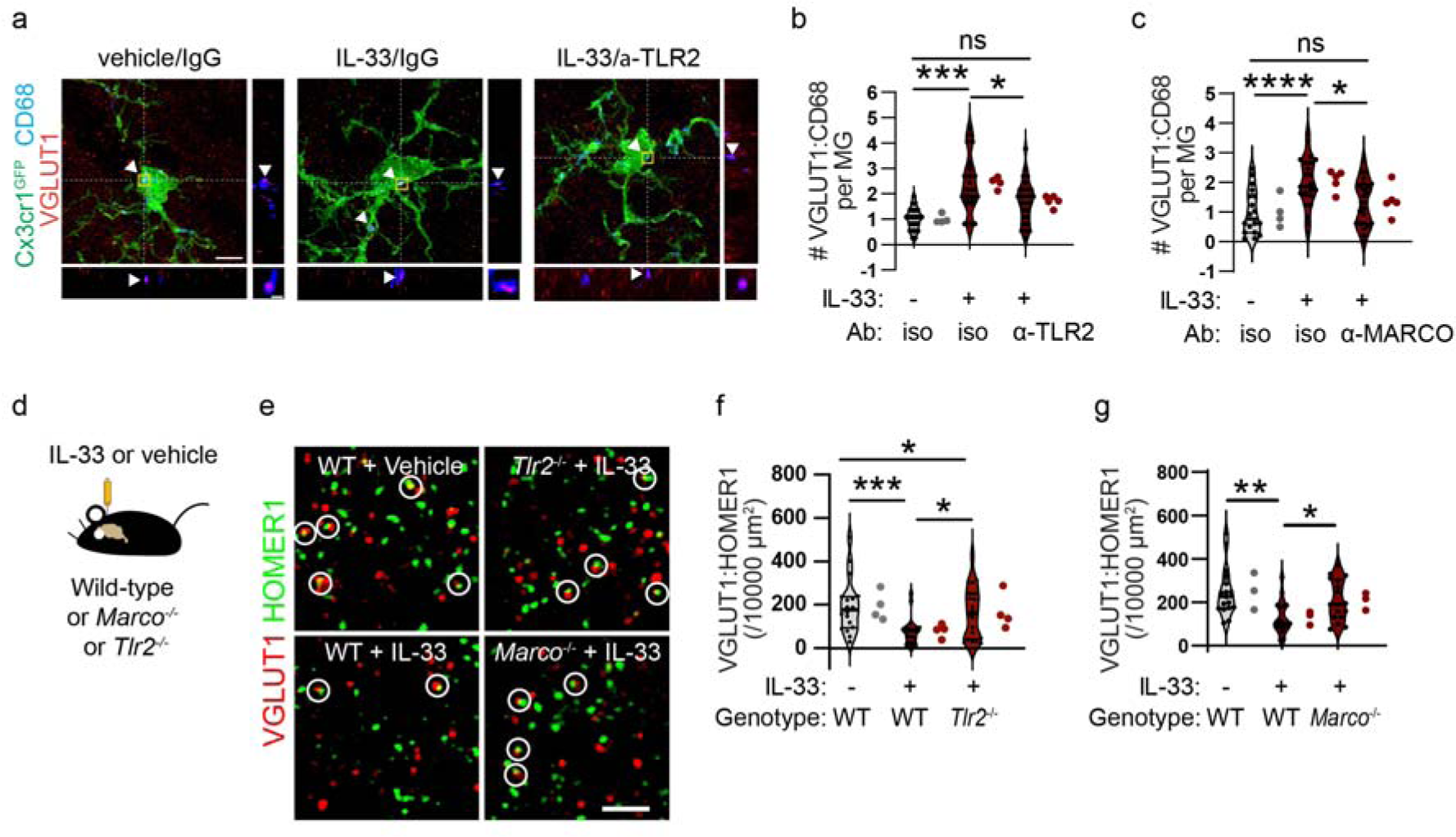
MARCO and TLR2 promote IL-33 dependent synapse engulfment and restrict excitatory synapse numbers. **a)** Representative images of Z-stack maximum projection from Cx3cr1^GFP^ microglia in the somatosensory thalamus for the indicated conditions. Arrowheads and orthogonal projections highlight engulfed VGLUT1 within CD68+ phagolysosomes. Vehicle=PBS. Scale bar: 5 µm (main panel) and 0.5 µm (inset). **b)** Quantification of VGLUT1 within CD68+ phagolysosomes within individual microglia after vehicle or IL-33 injection in the presence of TLR2 blocking antibody or isotype control (values normalized to vehicle+isotype control condition; n=24 microglia from 4 mice for vehicle+isotype control, n= 21 microglia from 4 mice for IL-33+isotype control, and n= 27 microglia from 5 mice for IL33+TLR2 blocking antibody). **c)** Quantification of VGLUT1 within CD68+ phagolysosomes within microglia after vehicle or IL-33 injection after co-injection of a MARCO blocking antibody or isotype control (values normalized to vehicle+isotype control condition; n=18 microglia from 4 mice for vehicle+isotype control, n= 23 microglia from 5 mice for IL-33+isotype control, and n= 21 microglia from 5 mice for IL33+MARCO blocking antibody). **d)** Schematic of intracerebroventricular injection of IL-33 in wild type, *Marco^-/^*^-^, or *Tlr2^-/-^* animals. **e)** Representative images of corticothalamic excitatory synapses defined by pseudocolocalization of pre- and post- synaptic proteins VGLUT1 and HOMER1 respectively for indicated conditions. White circles indicate colocalized puncta defined as presumptive functional synapses. Scale bar: 2 µm. **f)** Quantification of corticothalamic excitatory synapses in somatosensory thalamus after vehicle or IL-33 injection into *Tlr2^-/-^* animals or wild-type animals (n = 18 fields of view for wild-type+vehicle, and 20 fields of view for wild-type+IL-33 and *Tlr2^-/-^* + IL-33, 4 mice/condition). **g)** Quantification of corticothalamic excitatory synapses in somatosensory thalamus after vehicle or IL-33 injection into *Marco^-/^*^-^ or wild-type animals (for each condition n = 17 fields of view from 3 mice/condition). Data represented as median ± interquartile range for violin plots. Larger dots to the right of violin plots represent the average per individual mouse within that group. Mice from P15-P17 were used for all experiments. One-way ANOVA followed by post hoc Tukey’s comparison was used for all analysis. * *p*<0.05, ** *p*<0.01, *** *p*<0.001, **** *p*<0.0001. ***See also Figure S3.***

To determine whether MARCO or TLR2 signaling impacted synapse numbers, we quantified excitatory synapses after IL-33 exposure^13^ in genetic loss-of function models (**Fig. 2d;** model validation in **Fig. S3a**). We found a 2-fold reduction in functional excitatory synapses after IL-33 injection, as assessed by pseudocolocalization of pre- and postsynaptic proteins VGLUT1 and HOMER1^43^, consistent with our prior findings^13^. This depletion was significantly attenuated on either a *Tlr2* or a *Marco* deficient background in both thalamus (**Fig. 2e-g**) and cortex (**Fig. S3b-c**). Quantification of excitatory synapses after acute loss of function with α-MARCO or α -TLR2 antibodies phenocopied this effect, with somewhat more robust results (**Fig. S3d-e**). Taken together, these data indicate that both *Tlr2* and *Marco* are IL-33 targets that partly mediate its ability to promote synaptic protein engulfment and restrict synapse numbers.

### IL-33 promotes AP-1/FOS activation to drive target gene expression in microglia

To define the regulatory mechanisms responsible for IL-33-dependent gene expression programs, we next characterized the epigenetic changes in microglia responding to IL-33 (**Fig. 3A)**. We isolated purified microglia after i.c.v. injection of 500 ng IL-33 or vehicle (gating strategy in **Fig. S1A)**. We assessed chromatin accessibility using an assay for transposase-accessible chromatin sequencing (ATAC-seq; Buenrostro et al., 2013). We also performed chromatin immunoprecipitation sequencing for acetylation of histone H3 lysine 27 (H3K27ac ChIP-seq) to determine active regulatory regions. These data were cross-correlated with bulk RNA sequencing performed in parallel. Bulk RNA sequencing was highly consistent with findings obtained by single cell RNA sequencing **(Fig. S4A- B; Supplemental Table 2**). Representative examples of ATAC-seq and H3K27ac peaks from IL-33 vs. vehicle exposed microglia are shown in the vicinity of *Marco* and *Tlr2* (**Fig. 3B,** quality control in **Fig. S4C-D),** as well as quantification of total mRNA levels (**Fig. 3C**). Both reveal increased accessibility at promoter regions (yellow shading) as well as novel enhancers (purple shading) that were specifically induced in response to IL- 33.

**Figure 3:**
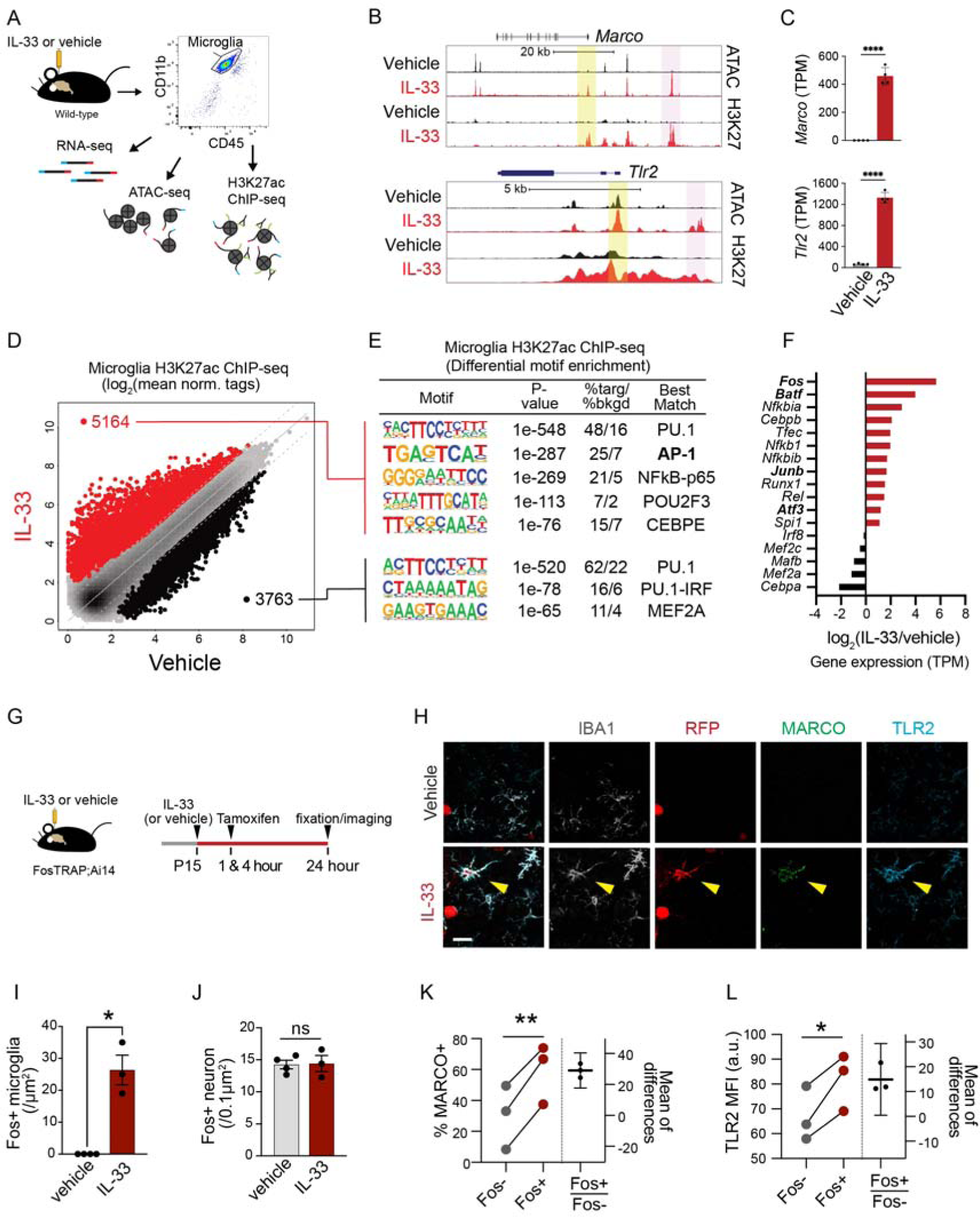
IL-33 promotes AP-1/Fos activation to drive target gene expression in microglia. **A)** Schematic of bulk RNA-seq, ATAC-seq and H3K27ac ChIP-seq paradigm. Vehicle=PBS. **B)** Browser tracks of ATAC-seq and H3K27ac ChIP-seq peaks in the vicinity of *Marco* and *Tlr2*. Yellow shading denotes promoter regions, pink shading denotes distal gene regulatory elements (enhancers). **C)** Bar graphs illustrate mRNA expression (transcripts per million, TPM) from bulk RNA-seq for *Marco* and *Tlr2*. Error bars = standard deviation. Each dot represents a mouse (two-tailed unpaired t-test). **D)** Scatter plot of normalized H3K27ac ChIP-seq in regions with ATAC-seq signal, in microglia after vehicle or IL-33 exposure. Data focuses on putative enhancers (chromatin regions > 3kb from transcriptional start site). Color codes indicate significant changes (FDR < 0.05 & FC >2) in H3K27ac ChIP-seq signal (IL-33 enriched= red, vehicle enriched=black). **E)** Enriched *de novo* motifs in distal open chromatin regions (enhancers) that gained or lost H3K27ac ChIP-seq signal after treatment with IL-33 or vehicle, showing best matched TFs binding to those motifs. **F)** Log_2_ fold-change of gene expression of all transcription factors that bind DNA elements identified in (**E**). All transcription factors shown have adj. p-value <0.001 by RNAseq. **G)** Experimental paradigm using the Fos-Trap2 allele crossed to the *Ai14* TdTomato reporter to label, or ‘trap’ Fos+ cells. Tamoxifen administered 1 and 4 hours after IL-33 or vehicle i.c.v. injection to capture Fos+ cells, animals sacrificed 20 hours after the last tamoxifen injection. **H)** Representative images of staining for Fos-TRAP (TdT+), MARCO and TLR2 after vehicle or IL-33 injection. Scale bar = 10 µm. **I)** Quantification of Fos+ microglia in the cortex after vehicle or IL-33 injection. Each dot represents a mouse (two-tailed unpaired t-test). **J)** Quantification of Fos+ neurons in the cortex after vehicle or IL-33 injection. Each dot represents a mouse (two-tailed unpaired t-test). **K)** Quantification of percent microglia that are MARCO+ comparing Fos- or Fos+ microglia in the cortex after IL-33 injection. Lines (left) connect paired measurements of Fos+ and Fos- microglia in the same mouse (two-tailed paired t-test). The right hand estimation plot shows the difference between the two means for each mouse, error bars indicate 95% confidence interval of that difference. **L)** Quantification of TLR2 mean fluorescent intensity from Fos- or Fos+ microglia in the cortex after IL-33 injection. Lines (left) connect paired measurements of Fos+ and Fos- microglia in the same mouse (two- tailed paired t-test). The right hand estimation plot shows the difference between the two means for each mouse, error bars indicate 95% confidence interval of that difference. Data represented as mean ± SEM for bar graphs. Mice at P30 were used for A-F. \**p*<0.05, \*\**p*<0.01, \*\*\*\**p*<0.0001. ***See also Figure S4.***

To globally examine this epigenetic signature and to determine potential transcriptional regulators of the IL-33 dependent response, we performed genome wide comparisons to identify chromatin regions that were open and active, as defined by both ATAC-seq and H3K27ac peaks. All peaks are listed in in Supplemental Table 3 and can be visualized in the UCSC data browser (https://genome.ucsc.edu/s/jschlachetzki/IL33_Microglia_mm10). We found a robust induction of *de novo* enhancers peaks in response to IL-33 (**Fig. 3D; Fig. S4E**). Motif enrichment analysis of regions that gained active open chromatin in response to IL-33 showed a significant enrichment for binding sites of adaptive-response type transcription factors (TFs), including AP-1 and NF-κB-p65 (**Fig. 3E; Fig. S4F**)^45, 46^. We also found suppression of MEF2, a TF associated with a microglial physiology/surveillance phenotype ^9, 47^. The myeloid lineage-defining pioneer factor PU.1 is required for chromatin opening, enabling subsequent accessibility of state-dependent TFs ^48^. PU.1 both gained and lost accessibility sites in response to IL-33, suggesting changes in its binding, but not necessarily that these changes are driving gene expression differences. We conclude that IL-33 markedly increased accessibility to stimulus-responsive transcription factors, including the AP-1 transcription factor (TF) complex.

AP-1 is a heterodimeric TF complex that includes members of four families: Fos, Jun, ATF/CREB, and Maf^49^. IL-33 exposed microglia had significantly upregulated gene expression of TFs in the AP-1 family, including *Fos*, *Batf, Junb,* and *Atf3 (***Fig. 3F**). We observed that *Fos* was the top TF induced after IL-33 at the transcriptional level, and that in myeloid cells, FOS binds to the de novo enhancers recruited by IL-33 in *Marco* and *Tlr2* (**Fig. S4G-H**)^50^. *Fos* is an immediate early gene whose induction is often used as a marker of neuronal activation. Its role in microglia is unknown, raising the question of whether FOS, as part of the AP-1 complex, is a regulator of IL-33 dependent gene expression.

We used a Fos-TRAP approach to permanently label cells that recently expressed *Fos*, using the *FosTrap2* knockin mouse crossed to a lox-stop-lox fluorescent reporter^51^. In this system, Fos induction drives expression of tamoxifen-inducible Cre recombinase. Administering tamoxifen then leads to excision of a stop cassette and permanently labels cells that expressed *Fos* after the tamoxifen pulse (**Fig. 3G**). Of note, while this allele is specific to *Fos* and two-fold more sensitive than TRAP1, it likely does not label all Fos+ cells. We found that IL-33 significantly increased the number of Fos-trapped microglia (**Fig. 3H-I**). *Fos* induction in microglia required expression of IL-33R on myeloid cells (**Fig. S4I**). In contrast, the number of Fos-trapped neurons was unchanged, suggesting no effect of IL-33 on neuronal activation (**Fig. 3J**). We found that *Fos*-trapped microglia in IL-33 exposed brains expressed significantly more TLR2 and MARCO protein relative to non-trapped microglia in the same sections (**Fig. 3K-L**). Taken together, our data suggest that IL-33 exposes AP-1 regulatory regions and promotes FOS/AP-1-mediated target gene expression.

### CNS-derived IL-33 acts through myeloid cells to restrict excitatory thalamic synapse numbers

We next investigated whether CNS-specific loss of IL-33 or myeloid-specific deletion of the IL-33 receptor impacted synaptic abundance and synaptic function. We deleted IL-33 from all CNS cells but not peripheral tissues (‘IL-33 cKO’) using the *hGFAPcre* transgenic line, which expresses Cre recombinase in neurons and glia in the forebrain and most astrocytes in the spinal cord (*hGFAPcre:Il33^fl/fl^* vs. *Il33^fl/fl^*, **Fig. S5a**)^52, 53^. Virtually all IL-33+ cells in the thalamus were astrocytes, and the remainder were oligodendrocytes (0% at P15 and 5.1% at P30, **Fig. S5b-c)**. In the somatosensory cortex, 80-90% of IL-33+ cells were astrocytes, and the remainder were oligodendrocytes (5.4% at P15 and 16.7% at P30; **Fig. S5d-e**), consistent with reports of oligodendrocyte expression in adult cortex (Gadani et al., 2015). Overall, IL-33 expression in the thalamus was more astrocyte-specific and robust than in cortex (**Fig. S5f**). In human tissues, IL-33 was robustly expressed in cortical astrocytes and detectable in some cortical neurons, consistent with our findings in mice (**Fig. S5g-h)**. In summary, during early postnatal development, most IL-33 expressing cells are astrocytes, and *hGFAPcre* efficiently deletes IL-33 in the brain.

We focused on the reciprocal connections between somatosensory cortex and thalamus, in part because the somatosensory thalamus is the first brain region to express IL-33 beginning at postnatal day 5^13^. This circuit is critical for somatosensory processing, but hyperexcitability in these connections is linked to type of childhood epilepsy known as absence epilepsy. The subtypes of synapses in the somatosensory thalamus include excitatory VGLUT2+ sensory afferents^54^, excitatory VGLUT1+ cortical afferents, and VGAT+ inhibitory synapses. All of these synaptic connections mature during a period of rapid IL-33 increase in thalamic astrocytes, accelerating after the second postnatal week^16, 17, 55^ (**Fig. S5i**).

We quantified synaptic subtypes at P30 by pseudocolocalization of pre and postsynaptic proteins^43^. We found a significant increase in the number of corticothalamic excitatory synapses (VGLUT1:Homer1) in IL-33 cKO animals vs. littermate controls (**Fig. 4a-b**). This was phenocopied with loss of myeloid IL-33R (*Cx3cr1^CreERT2+/-^:Il1rl1^fl/fl^ vs. Cx3cr1^CreERT2+/-^:Il1rl1^wt/wt^*; **Fig. 4c**). We also observed a more modest increase in brain stem to thalamic excitatory synapses in both genotypes (VGLUT2:Homer1; **Fig. 4d-f**). In contrast, the number of inhibitory synapses (VGAT:Gephyrin) was significantly decreased in both genotypes (**Fig. 4g-i**). We did not observe gross structural alterations in thalamic neurons of IL-33 cKO animals, as assessed by Scholl analysis of neuronal branching (**Fig. S5j-k**). There were similar but more modest changes in synapse numbers in somatosensory cortex of IL-33 cKO animals (**Fig. S5l-n**), consistent with the lower expression of IL-33 in cortex. In summary, CNS-specific loss of IL-33 resulted in excess excitatory synapses and fewer inhibitory synapses at both nodes of the corticothalamic circuit.

**Figure 4:**
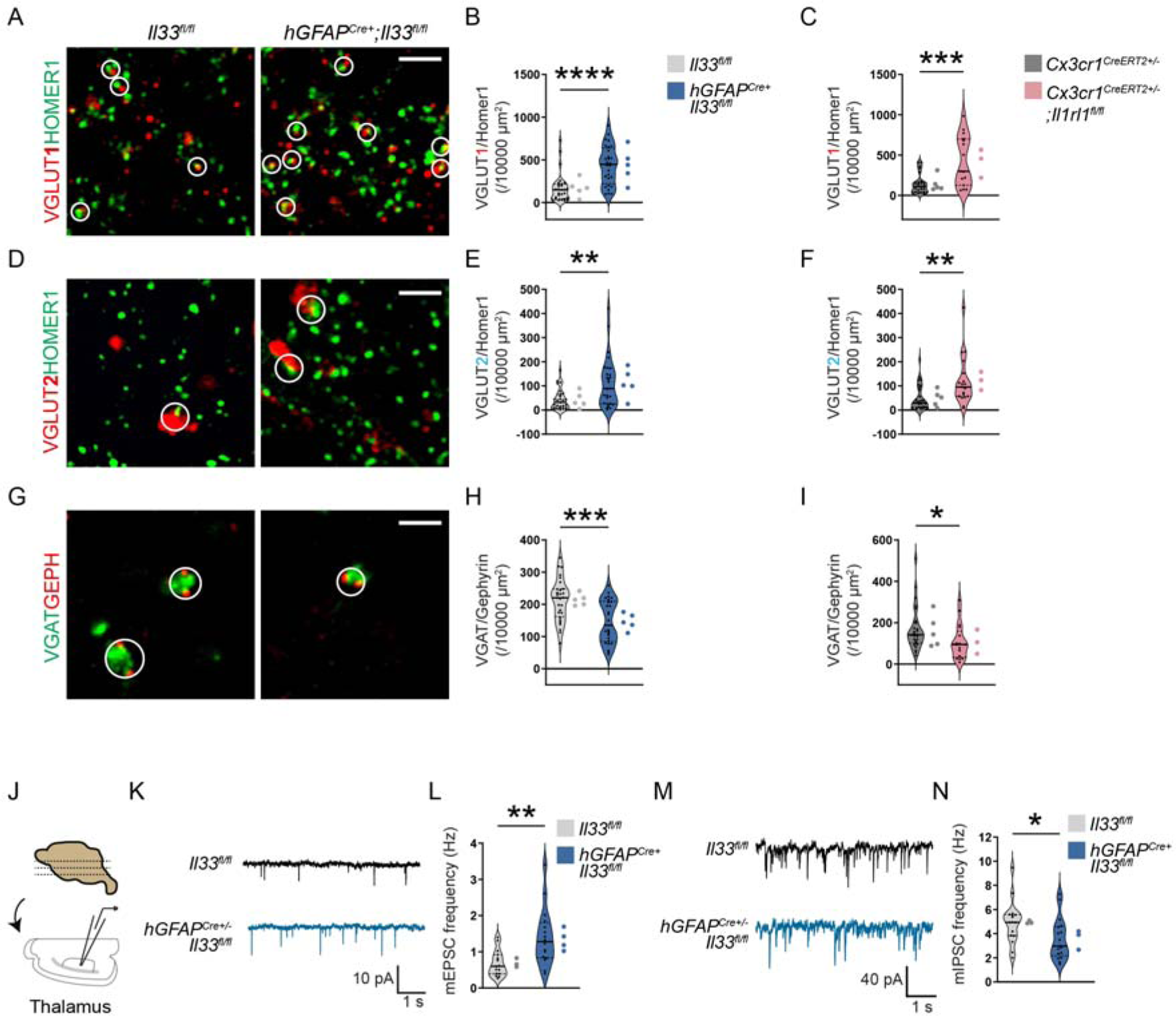
CNS-derived IL-33 acts through myeloid cells to restricts excitatory thalamic synapse numbers. **a)** Representative images of corticothalamic excitatory synapses as defined by colocalized presynaptic (VGLUT1) and postsynaptic(HOMER1) puncta in *hGFAPCre+;Il33^fl/fl^* vs. *Il33^fl/fl^* control. Circles indicate co-localization, defining a functional synapse. Scale bar = 2 µm. **b)** Quantification of corticothalamic excitatory synapses in *hGFAPCre+;Il33^fl/fl^* vs. *Il33^fl/fl^* control. n= 28 fields of view, 5 mice/genotype. **c)** Quantification of corticothalamic excitatory synapses with myeloid-specific deletion of IL-33 receptor (*Cx3cr1Cre^ERT2^*^+/-^;*Il1rl1^fl/fl^*) vs. control (*Cx3cr1^CreERT2^* ^+/-^). n= 15 fields of view from 3 mice in *Cx3cr1^CreERT2^*^+/-^;*Il1rl1^fl/fl^* and n= 25 fields of view from 5 mice in *Cx3cr1^CreERT2^*^+/-^. **d)** Representative images of brainstem afferent synapses as defined by colocalized pre- (VGLUT2) and post- (HOMER1) synaptic puncta in *hGFAPCre+;Il33^fl/fl^* vs. *Il33^fl/fl^*. Circles indicate co-localization, defining a functional synapse. Scale bar = 2 µm. **e)** Quantification of brainstem afferent synapses in *hGFAPCre+;Il33^fl/fl^* vs. *Il33^fl/fl^* control mice. n= 28 fields of view, 5 mice/genotype. **f)** Quantification of brainstem afferent synapses after myeloid-specific deletion of IL-33 receptor (*Cx3cr1Cre^ERT2^*^+/-^;*Il1rl1^fl/fl^*) vs. control (*Cx3cr1^CreERT2^* ^+/-^). *Cx3cr1^CreERT2^*^+/-^;*Il1rl1^fl/fl^* : n= 19 fields of view from 3 mice. *Cx3cr1Cre^ERT2^*^+/-^: n= 29 fields of view from 5 mice. **g)** Representative images of thalamic inhibitory synapses as defined by colocalized presynaptic (VGAT) and postsynaptic (Gephyrin) puncta in *hGFAPCre+;Il33^fl/fl^* vs. *Il33^fl/fl^* control. Circles indicate co- localization, defining a functional synapse. Scale bar = 2 µm. **h)** Quantification of thalamic inhibitory synapses in *hGFAPCre+;Il33^fl/fl^* vs. *Il33^fl/fl^* control. n= 28 fields of view, 5 mice/genotype. **i)** Quantification of thalamic inhibitory synapses in myeloid-specific deletion of IL-33 receptor (*Cx3cr1Cre^ERT2^*^+/-^;*Il1rl1^fl/fl^*) vs. control (*Cx3cr1^CreERT2^*^+/-^). *Cx3cr1^CreERT2^*^+/-^;*Il1rl1^fl/fl^* : n= 17 fields of view from 3 mice. *Cx3cr1^CreERT2^*^+/-^: n= 26 fields of view from 5 mice. **j)** Experimental paradigm for whole cell patch-clamp electrophysiology of somatosensory thalamic neurons. **k)** Representative traces of miniature excitatory post synaptic currents (mEPSC) from somatosensory thalamus over a 5 second recording period. **l)** Quantification of mEPSC frequency in somatosensory thalamic neurons (*Il33^fl/fl^* control: n=17 neurons from 3 mice, *hGFAPCre+;Il33^fl/fl^* : n=18 neurons from 4 mice, 2 independent experiments, all mice age P26-P33). **m)** Representative traces of miniature inhibitory post synaptic currents (mIPSC) from somatosensory thalamus over a 5 second recording period. **n)** Quantification of mIPSC frequency in somatosensory thalamic neurons (n=16 neurons from 3 mice in *Il33^fl/fl^*, n=17 neurons from 3 mice in *hGFAPCre+;Il33^fl/fl^*, 2 independent experiments, all mice age P28- P36). Data represented as median ± interquartile range for violin plots. Larger dots to the right of violin plots represent the average per individual mouse within that group. Two-tailed unpaired t-test used for all analyses. P28-P30 mice were used for b, e, and h. P32-P35 mice were used for c, e, and i. * *p*<0.05, ** *p*<0.01, *** *p*<0.001, **** *p*<0.0001. ***See also Figure S5.***

To determine whether these changes in synapse numbers were also associated with changes in synaptic function, we performed whole-cell patch clamp electrophysiology in somatosensory thalamus of IL-33cKO mice (**Fig. 4j**). Miniature excitatory post synaptic currents (mEPSC) were higher in frequency in IL-33cKO mice (**Fig. 4k-l**). This was consistent with an increase in synapse numbers that was similar in magnitude to what we observed with immunostaining. However, mEPSC amplitude was unchanged**Table S4**). In addition, miniature inhibitory postsynaptic currents (mIPSCs) were reduced in frequency in IL-33 cKO (**Fig. 4m-n**), consistent with fewer inhibitory synapses, but unchanged in amplitude (**Table S4**). Taken together, these studies suggest that CNS- derived IL-33 acting on myeloid cells increases excitatory/inhibitory ratio by both restricting excitatory synapse numbers and promoting inhibitory synapse numbers, without altering synaptic strength.

### IL-33-IL1RL1 signaling limits seizure susceptibility

Thalamic hyperexcitability can lead to seizures in rodent models^18, 56^ and is particularly associated with childhood absence epilepsy^57^. To investigate whether the increased excitation after loss of CNS-derived IL-33 impacted brain electrical activity, we performed electrocorticography (ECoG) recordings. Leads were placed in primary somatosensory cortex (S1) and prefrontal (PFC) cortex in freely behaving juvenile mice (5-6 weeks old; **Fig. 5a**). One hour of baseline recording in the home cage environment revealed the presence of brief (1-3 second) spontaneous spike-wave discharges (SWDs) in 6/10 IL-33 cKO mice but these were never observed in littermate controls (**Fig. 5b-c**). These SWDs were observed simultaneously in both cortical regions (S1 and PFC; **Fig. 5b**), indicating that they were bona fide events, and were frequently associated with behavioral arrest in simultaneous videorecording. SWD had a characteristic internal frequency of 4-6 Hz, with a harmonic peak at 12 Hz, clearly distinguishing them from sleep spindles (**Fig. 5d**). Taken together, this phenotype resembled typical absence type (‘petit mal’) seizures seen in various rodent models of childhood absence epilepsy, which results from alterations in corticothalamic function^57–59^.

**Figure 5:**
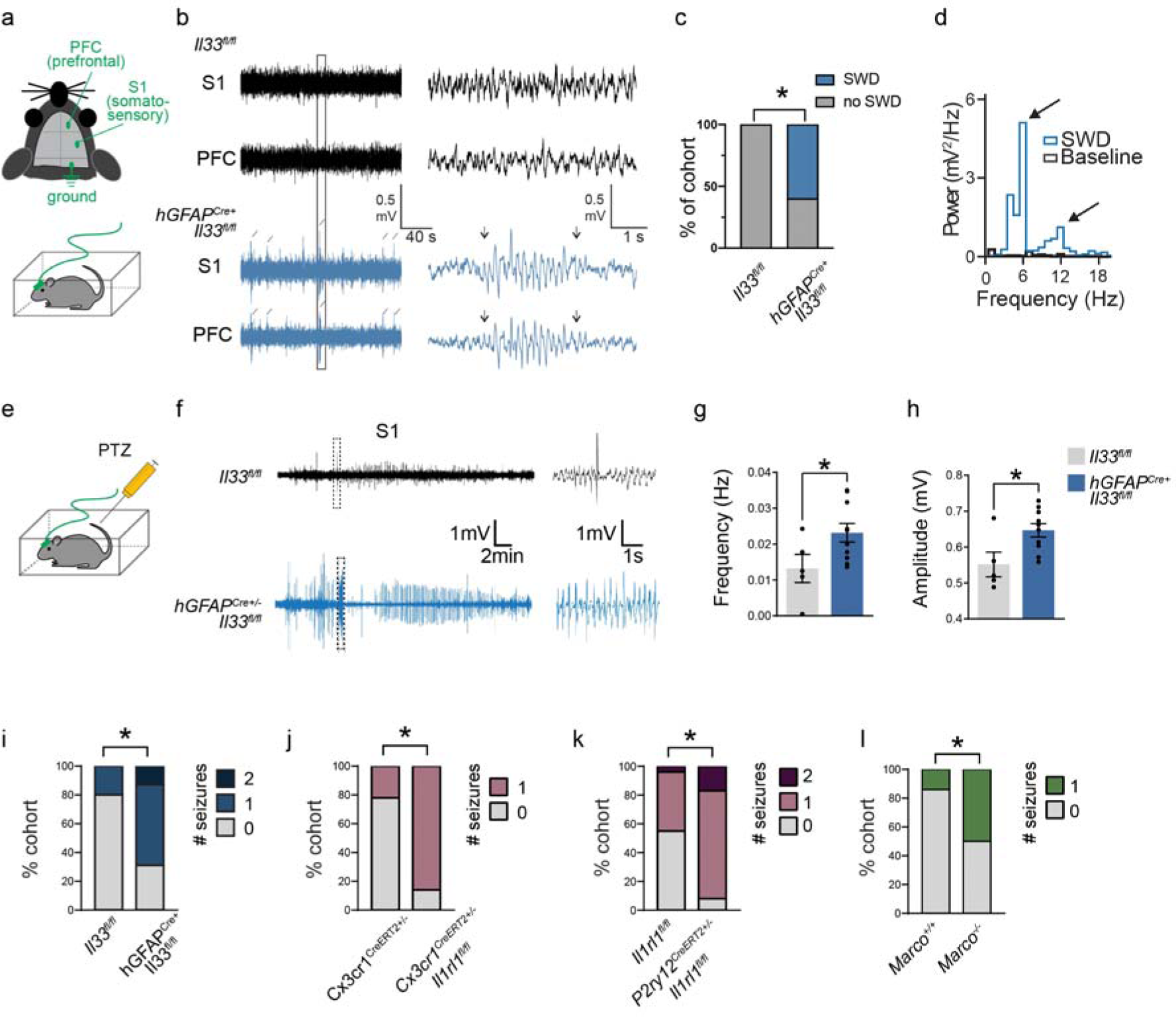
IL-33-IL1RL1 signaling limits seizure susceptibility. **a)** Schematic of lead placement and setup for electrocorticography (ECoG) in 5–6 week-old mice. **b)** Representative traces of recordings from somatosensory and prefrontal cortex of freely behaving mice. Left shows several minutes of recording with detected spiking events indicated by diagonal lines. Boxed area on the left indicates inset which is shown at a higher time scale on the right. The inset highlights a representative spike-wave discharge (SWD) lasting about 3 seconds, occurring in *hGFAP^Cre+^;Il33^fl/fl^*. Arrowheads indicate the beginning and end of the event. **c)** Quantification of percent of cohort with synchronized spike-wave-discharge (SWD) events in both cortical areas, collected during the one-hour recording session. n=10 *hGFAP^Cre+^;Il33^fl/fl^* mice and 5 *Il33^fl/fl^* mice (Fisher’s exact test). **d)** Representative fast Fourier transform of a spike wave discharge observed in *hGFAP^Cre+^;Il33^fl/fl^* mice, demonstrating characteristic peak at 6 Hz and harmonic peak at 12 Hz. **e)** Schematic of ECoG recordings after injection of the GABA-A receptor antagonist pentylenetetrazol (PTZ). Mice were recorded for one hour. **f)** Representative traces of recordings from somatosensory cortex of PTZ-injected mouse. The left trace shows several minutes of recording time. Boxed area indicates a spiking event shown at a higher time scale in the inset on the right. **g-h)** Quantification of total spike frequency **(G)** and average amplitude of detected spike events **(H)** from somatosensory cortex during one-hour recording session. n=10 *hGFAP^Cre+^;Il33^fl/fl^* mice and 5 *Il33^fl/fl^* mice (two-tailed unpaired t-tests). **i)** PTZ injection was followed by one hour of video recording, with behavioral scoring of observed seizure events on a Racine scale by a blinded observer. Quantification shows percent of cohort that experienced generalized tonic-clonic seizures during the recording. n=16 *hGFAPCre^+^;Il33^fl/fl^* mice and 15 *Il33^fl/fl^* mice from 4 independent experiments. Age P29-P35, Fisher’s exact test. **j)** PTZ injection followed by behavioral scoring of seizure events, showing percent of cohort that experienced generalized tonic-clonic seizures during the recording. n=7 *Cx3cr1^CreERT2^*^+/-^;*Il1rl1^fl/fl^* mice and n=9 *Cx3cr1^CreERT2^*^+/-^ mice from 3 independent experiments. Age P29-P35, Fisher’s exact test. k) PTZ injection followed by behavioral scoring of seizure events, showing percent of cohort that experienced generalized tonic-clonic seizures during the recording. n=12 *P2ry12CreERT2*+/-;*Il1rl1^fl/fl^* mice and n=22 *Il1rl1^fl/fl^* mice from 5 independent experiments. Age P29-P35, (Fisher’s exact test). l) PTZ injection followed by behavioral scoring of seizure events, showing percent of cohort that experienced generalized tonic-clonic seizures during the recording. n=21 wild-type and 16 *Marco*^-/-^ from 4 independent experiments (Fisher’s exact test). Data represented as mean ± SEM for bar graphs. Each dot represents mice. Mice from P35-40 were used for a-h. ECoG; electrocorticography, PTZ; pentylenetetrazol, S1; somatosensory cortex, PFC; prefrontal cortex. \**p*<0.05. ***See also Figure S6.***

To determine whether IL-33 cKO mice were also more susceptible to convulsive (‘grand mal’) seizures, we performed ECoG recordings in the same animals after injection of a chemoconvulsant, pentylenetetrazol (PTZ; 50 mg/kg), a GABA_A_ receptor blocker (**Fig. 5e**). We found an increase in the frequency and amplitude of spikes in response to PTZ in IL-33 cKO vs. littermate controls in both somatosensory and prefrontal cortex (**Fig. 5f-h; Fig S6a**). We also behaviorally quantified seizures in independent cohorts of mice that had not undergone lead placement by video recording for one hour after injection of PTZ, and blind scoring seizure events on a Racine scale. Consistent with our ECoG findings, we observed a significantly increased incidence of generalized tonic-clonic seizures in IL-33 cKO animals in response to PTZ: 11/16 IL-33 cKO mice had at least one seizure, compared to 3/15 littermate controls (**Fig. 5i**). Increased c-Fos staining of IL-33 cKO animals after PTZ was consistent with these findings (**Fig. S6b-d**). IL-33 cKO mice were also more susceptible to another chemoconvulsant, Kainic Acid (KA), a glutamate receptor agonist (**Fig. S6e-f**), further supporting their increased seizure susceptibility.

Importantly, this increase in seizure susceptibility was phenocopied after PTZ injection in mice with conditional deletion of IL-33R in myeloid cells using Cx3cr1^CreERT2^ (**Fig. 5J**; 6/7 *Cx3cr1^CreERT2+/-^:Il1rl1^fl/fl^* vs. 2/9 littermate *Cx3cr1^CreERT2+/-^* mice had at least one seizure) or microglia specific deletion of IL-33R using P2ry12 ^CreERT2^ (**Fig. 5k**; 11/12 *P2ry12^CreERT2+/-^:Il1rl1^fl/fl^* vs. 10/22 littermate *Il1rl1^fl/fl^* mice had at least one seizure). Moreover, seizure susceptibility was increased in juvenile *Marco* deficiency mice (**Fig 5i**; 8/16 *Marco* deficiency vs. 3/21 littermate wild-type mice had at least one seizure). However, *Tlr2* deficiency was not sufficient to increase seizure susceptibility in juvenile mice (**Fig. S6g-h**). Taken together, these results indicate that IL-33-Il1rl1 signaling - operating locally in the CNS and acting through myeloid cells – coordinates functional responses in microglia that restrict spontaneous epileptiform activities and limits seizure susceptibility.

## Discussion

IL-33 is a tissue resident cytokine initially defined as a regulator of allergic immune responses, but increasingly implicated in development, homeostasis, and remodeling^60^. Tissue resident macrophages, including microglia, are essential mediators of remodeling but their roles as direct targets of IL-33 remain underexplored. Our data reveal a critical and targeted role for IL-33, acting through microglia, in shaping the microglial epigenomic landscape (**Fig. 6.)** We demonstrate that IL-33 renders target genes accessible to stimulus-responsive TFs and coordinates a phagocytic gene expression program. This is consistent with a series of studies demonstrating that microglia are highly responsive to contextual cues^7, 10, 11, 61, 62^, and suggests that IL-33 is one factor mediating this context sensitivity.

**Figure 6:**
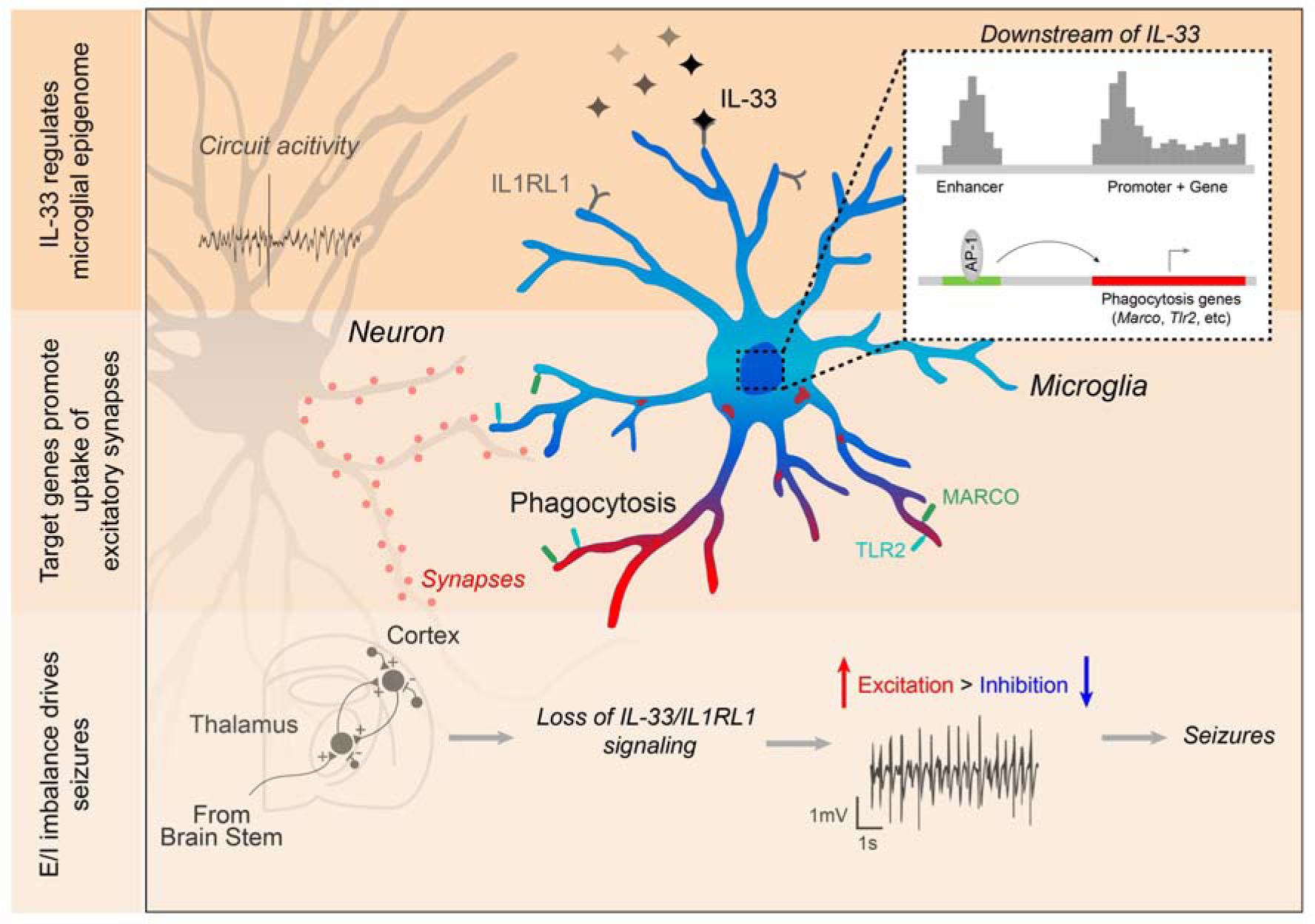
Graphical abstract. • IL-33 promotes state-dependent transcription factor expression in microglia and induces gene expression programs associated with sensing and scavenging. • The scavenger receptor MARCO and the pattern recognition receptor TLR2 are two downstream target genes that partly mediate IL-33’s effects on microglial uptake of synaptic proteins • Loss of CNS-derived IL-33 or its receptor on myeloid cells leads to corticothalamic excitability by altering synapse numbers and increases seizure susceptibility.

We demonstrate that IL-33 impacts synapse numbers rather than synaptic strength, consistent with a role in synapse remodeling. This brain-specific form of tissue remodeling is required for optimal neural circuit function by regulating the balance of excitatory to inhibitory synapses. Although neuronal activity is one signal that modulates IL-33 dependent engulfment, its impact is to promote microglial phagocytic function rather than acute feedback control of neuronal activity as shown in other contexts^63^. Our data demonstrate that loss of IL-33 signaling leads to hyperexcitability in a core circuit implicated in seizure generation. We identify MARCO and TLR2 as two top functional candidates downstream of IL-33 signaling, however, *Tlr2* deficiency alone is not sufficient to phenocopy the functional impact of IL-33 on seizure susceptibility. This could be due to homeostatic compensation. Taken together, these data suggest that IL-33 coordinates a broader ‘sensing and scavenging’ phagocytic program that is distinct from previously described mechanisms of microglial synaptic engulfment.

Neural circuit hyperexcitability is a phenotype that has been implicated in multiple neurodevelopmental disorders including epilepsy, autism, and schizophrenia^64^. Neuroimmune dysfunction is also increasingly implicated in these pathologies, although mechanistic evidence is lacking^4^. Microglia are attractive potential therapeutic targets in epilepsy^65^, particularly with the emergence of immunotherapies. Our study defines a core pathway regulating microglial phagocytic function, suggesting potential avenues to towards immune-mediated therapies for epilepsy and other neurodevelopmental disorders.

## Acknowledgements

We are grateful to members of the Molofsky Lab for helpful comments on the manuscript, to Dr. Mercedes Paredes for pathology expertise, and to Irene Lew for help with animal husbandry. Thanks to the UCSF Laboratory for Cell Analysis and Center for Advanced technology for technical contributions. Imaging was performed at the Gladstone Institutes’ Histology & Light Microscopy Core.

## Funding

A.V.M is supported by the Pew Charitable Trusts, the Brain and Behavior Research Foundation, NIMH (R01MH119349, MH125000, and DP2MH116507), and the Burroughs Welcome Fund. This study was supported in part by the HDFCCC Laboratory for cell Analysis Shared Resource facility through a shared grant from NIH (P30CA082103). J.T.P is supported by NIH & NINDS R01 NS096369-01, DoD EP150038, NSF 1608236, and the Gladstone Institutes Animal Facility Grant RR18928. F.S.C. is supported by NINDS NS111819.

## Authors contributions

Conceptualization: R.T.H., I.D.V., A.B.M., J.T.P., and A.V.M. Methodology: R.T.H, I.D.V., F.S.C., L.C.D, and T. J; Investigation: R.T.H., I.D.V., F.S.C., J. C. M. S., A.J., E.A., J.T.B. and H N-I; Writing – Original Draft: R.T.H., I.D.V., and A.V.M.; Writing – Review & Editing: all co-authors; Funding Acquisition: A.V.M. Resources: A.V.M. and J.T.P. Supervision, A.V.M., C.K.G. and J.T.P.

## Declaration of interests

The authors declare no competing interests.

## Data and materials availability

Supplement contains additional data. All data needed to evaluate the conclusions in the paper are present in the paper or the Supplementary Materials. Bulk RNA, ATAC, H3K27ac ChIP-sequencing and scRNAseq data of microglia post i.c.v. injection of IL-33 or vehicle are available through GEO [number pending], and epigenomic data has been uploaded to the UCSC data browser (https://genome.ucsc.edu/s/jschlachetzki/IL33_Microglia_mm10).

## Supplemental Figures

**Figure S1:**
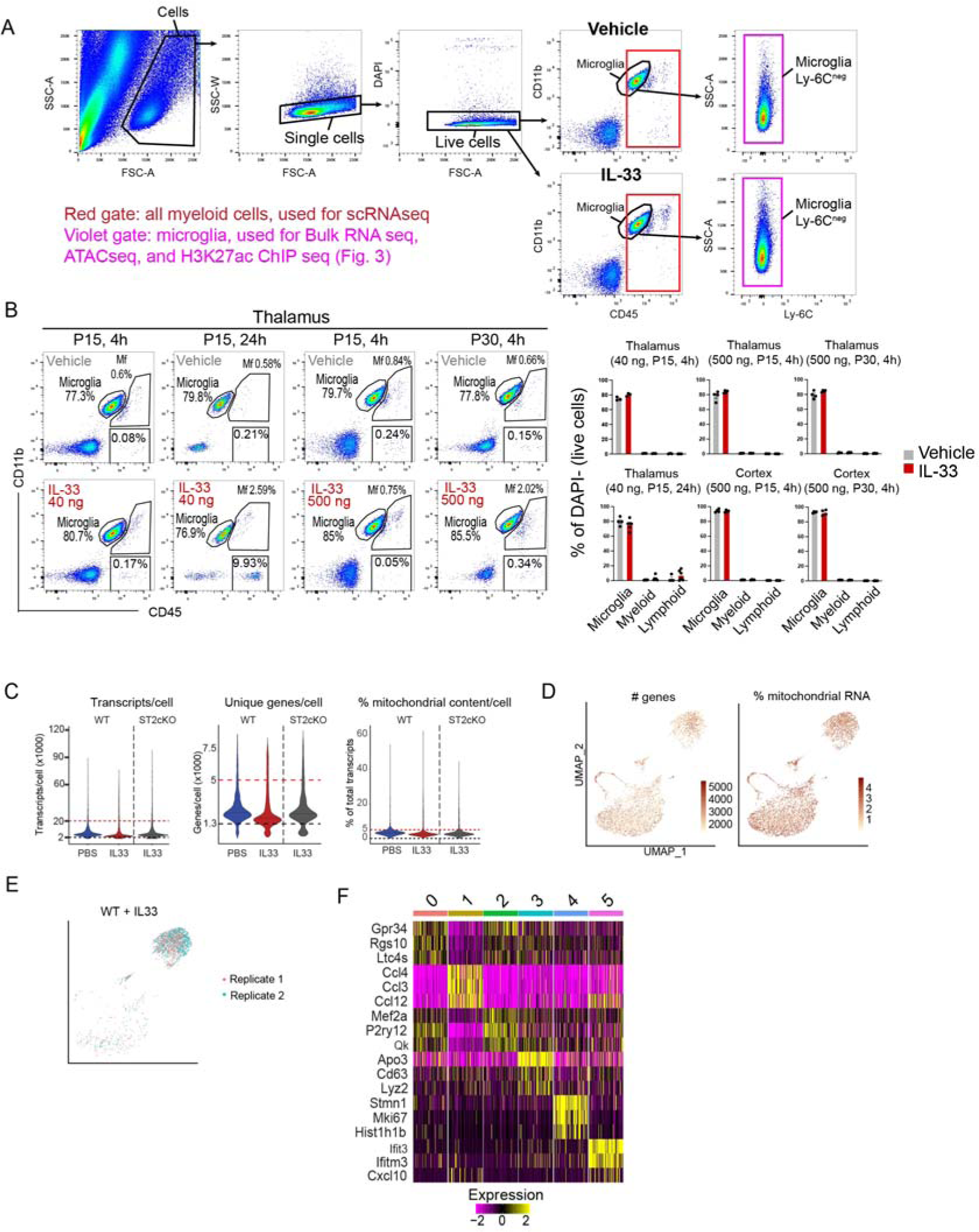
Quality control and additional data defining the microglial response to IL-33, related to Figure 1. **a)** Gating strategy for isolation by FACS of all CD45+ (red) for scRNAseq in Figure 1, or microglia only (CD11b+, CD45+, Ly-6C-, purple) for bulk RNA/ATAC/ChIP seq (Figure 3). **b)** Gating strategy and percentage of microglia, myeloid and lymphoid cells after vehicle or IL-33 as gated by CD11b and CD45. Representative plots are from Thalamus, quantifications on the right include Thalamus and Cortex. Each dot represents a mouse. **c)** Violin plots of scRNAseq data showing transcripts/cell, unique genes/cell and % mitochondrial content/cell for each sample. Cut off boundaries are marked (upper: dotted red line, bottom: dotted black line). **d)** Feature plots for number of genes and % mitochondrial RNA for all samples combined from scRNAseq data. Each dot represents a cell. **e)** Feature plot showing correlation between replicates for WT + IL-33 scRNAseq sample. **f)** Heatmap for the top 3 genes in each cluster from scRNAseq data.

**Figure S2:**
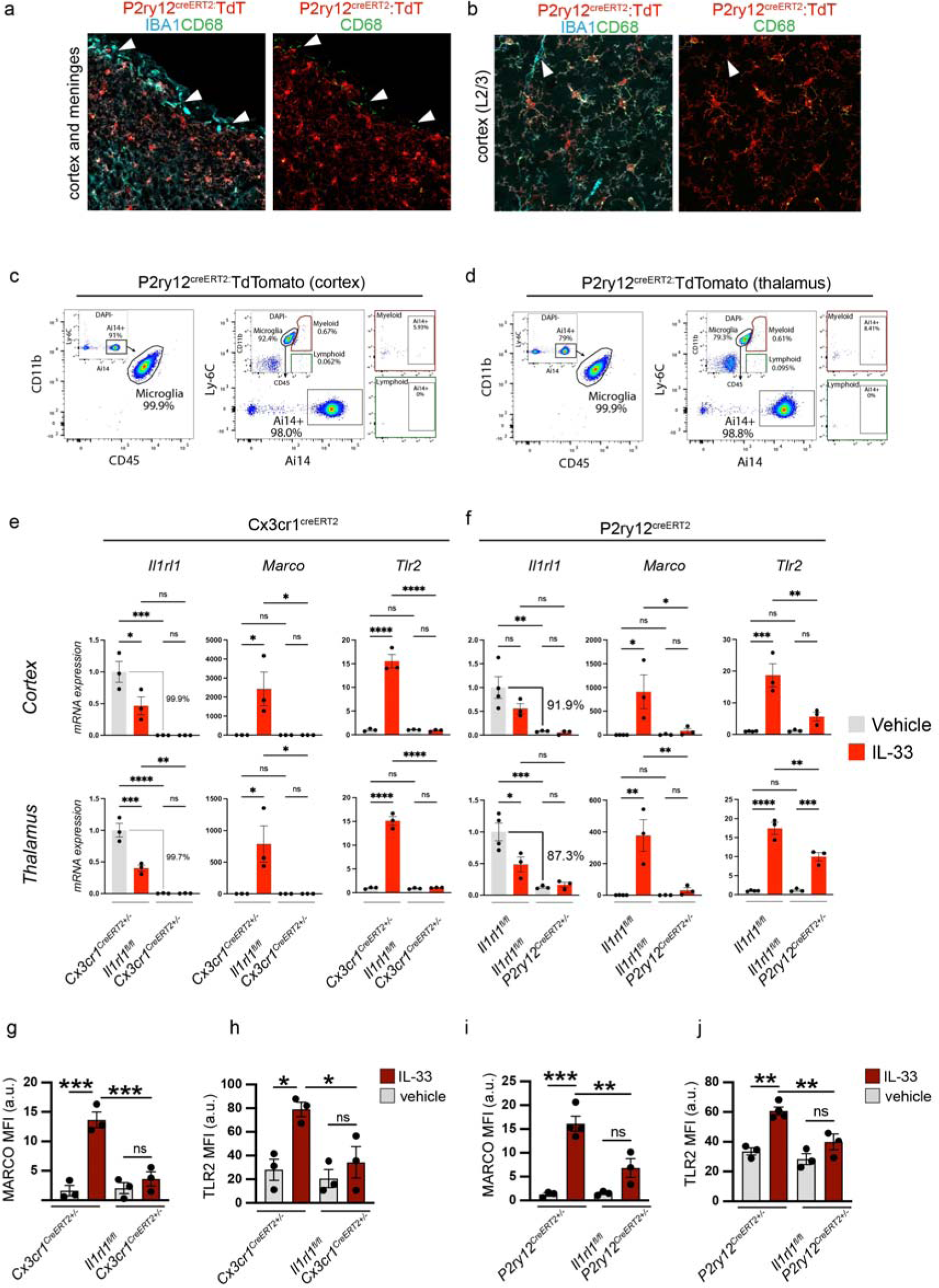
Additional data defining the microglial response to IL-33, related to Figure 1. **a-b)** Representative images for P2ry12^creERT2^ crossed to a R26R-TdTomato reporter (*Ai14*). Staining for TdT, IBA1 and CD68. Left panel (G) at lower power shows cortex with overlying meninges. Arrowheads indicate meningeal macrophages that are CD68+IBA1+TdT^neg^. Right panels **(H)** shows L2/3 cortex. Arrowheads indicate perivascular macrophage that is CD68+IBA1+TdT^neg^. **c-d)** Representative flow cytometry plots of Cortex **(I)** and Thalamus **(J)** showing P2ry12^creERT2^ driven Ai14 (TdTomato) expression. Two gating strategies shown: Left panels show gating on Ai14+ followed by CD11b and CD45. Right panels show gating on microglia, myeloid and lymphoid based on CD11b and CD45 followed by Ly-6C and Ai14. **e-f)** qRT-PCR of *Il1rl1*, *Marco* and *Tlr2* expression in cortical and thalamic microglia from Cx3cr1^creERT2^ **(e)** and P2ry12^creERT2^ **(f)** mice. Values were normalized to housekeeper (*Hmbs*) and control + vehicle (PBS) condition. In Cx3cr1^creERT2^ control=Cx3cr1^creERT2+/-^. In P2ry12^creERT2^ control=*Il1rl1^fl/fl^.* Each dot represents a mouse. Two-way ANOVA followed by Tukey’s post hoc comparison (genotype and treatment). **g-j)** Mean fluorescence intensity for MARCO and TLR2 protein in cortex from Cx3cr1^creERT2^ **(g-h)** and P2ry12-^creERT2^ **(i-j)** mice. Each dot represents a mouse. Two-way ANOVA followed by Tukey’s post hoc comparison (genotype and treatment). Data represented as mean ± SEM for bar graphs. \**p*<0.05, \*\**p*<0.01, \*\*\**p*<0.001, \*\*\*\**p*<0.0001.

**Figure S3:**
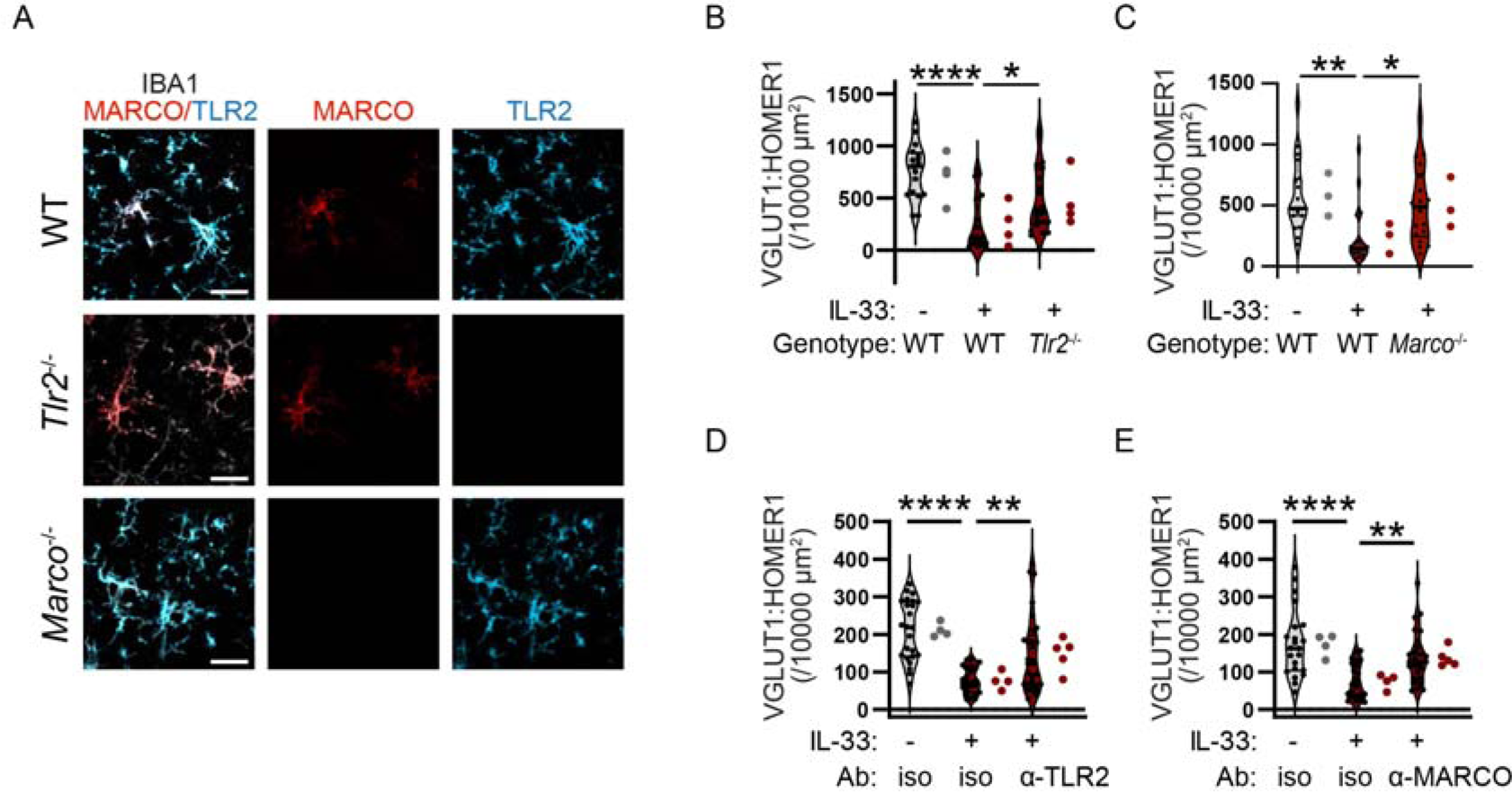
Validation of TLR2 and MARCO deficient animals and impact on cortical synapse numbers, related to Figure 2. **a)** Representative images of MARCO and TLR2 immunostaining in wild-type (top), *Tlr2* (middle), or *Marco* (bottom) deficiency animals 18 hours after 40 ng of IL-33 i.c.v. injection at P17. Scale bar = 20 µm. **b)** Quantification of intracortical synapses in somatosensory cortex after vehicle or IL-33 injection into *Tlr2^-/-^* animals or wild-type animals (n=17 fields of view for wild-type+vehicle, n=19 fields of view for wild-type+IL-33, n=20 fields of view for *Tlr2^-/-^* + IL-33, 4 mice/condition). **c)** Quantification of intracortical synapses in somatosensory cortex after vehicle or IL-33 injection into *Marco*^-/-^ animals or wild-type animals (n=19 fields of view for wild-type+vehicle, n=18 fields of view for wild-type+IL-33, n=18 for *Marco*^-/-^ + IL-33, 3 mice/condition). **d)** Quantification of corticothalamic synapses in somatosensory thalamus after vehicle or IL-33 injection in the presence of TLR2 blocking antibody or isotype control (n=24 fields of view from 4 mice for vehicle+isotype control and IL-33+isotype control, and n=29 fields of view from 5 mice for IL-33+ α- TLR2). **e)** Quantification of corticothalamic synapses in somatosensory thalamus after vehicle or IL-33 injection in the presence of MARCO blocking antibody or isotype control (n=22 fields of view from 4 mice for vehicle +isotype control, n=23 fields of view from 4 mice for IL-33+isotype control, and n=26 fields of view from 5 mice for IL-33+ α-MARCO). Data represented as median ± interquartile range for violin plots. Larger dots to the right of violin plots represent the average per individual mouse within that group. One-way ANOVA followed by post hoc Tukey’s comparison was used for all analysis. \**p*<0.05, \*\**p*<0.01, \*\*\*\**p*<0.0001.

**Figure S4:**
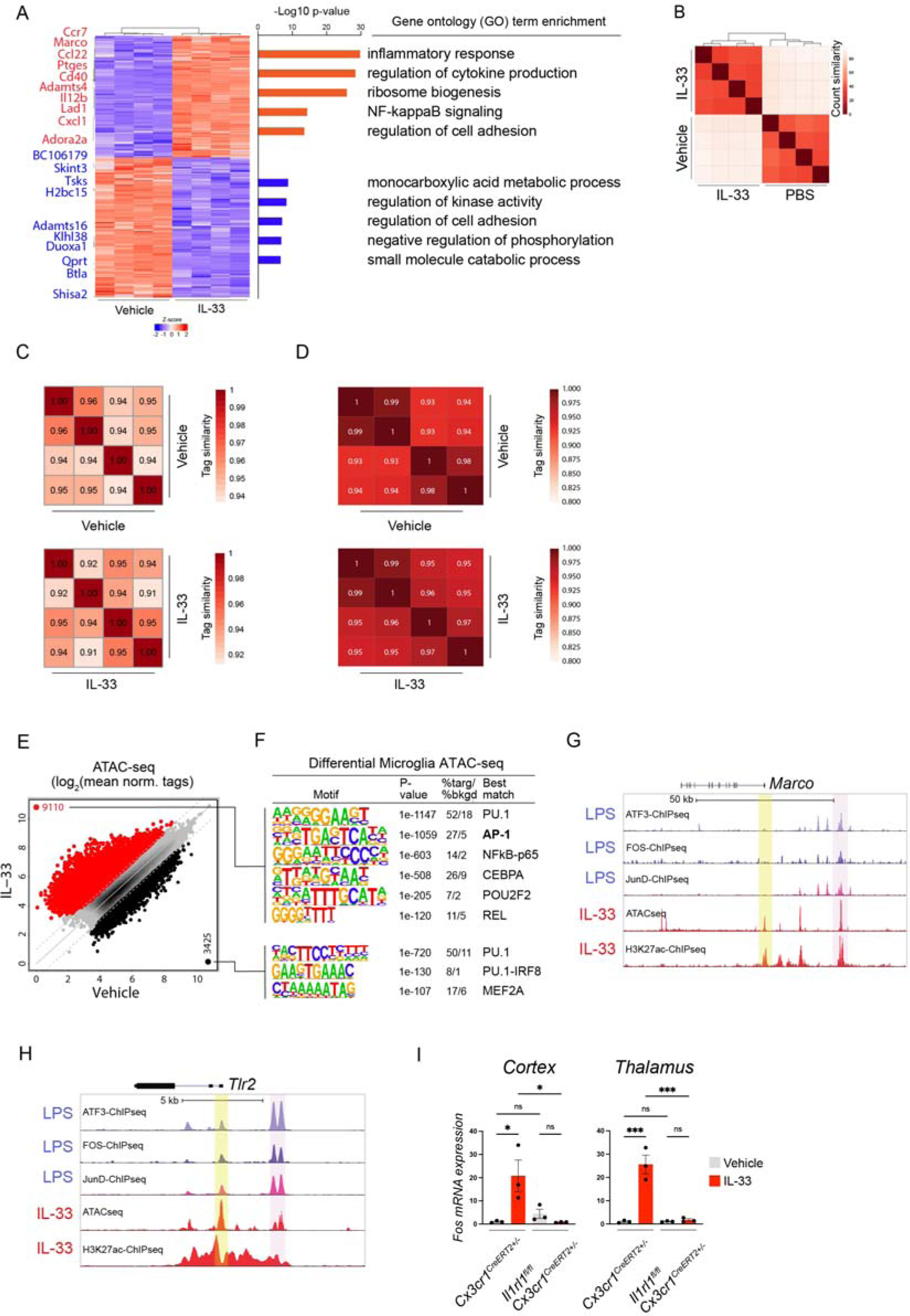
Transcriptomic and epigenomic profiling of microglia after IL-33 exposure, related to Figure 3. **A)** Heatmap of differentially expressed genes in cortical microglia four hours after vehicle (PBS) or 500 ng of IL-33 treatment (P_adj_ < 0.01). Left: Top 10 up (red) and downregulated (blue) genes indicated. Right: Top GO categories associated with differentially expressed genes (P_adj_ < 0.01, fold-change>2), upregulated (red) and downregulated (blue). **B)** Heatmap of sample-to-sample Pearson correlation of bulk RNA-seq replicates. **C)** Heatmaps of sample-to-sample correlation for ATAC-seq replicates. Values indicate Pearson correlation. **D)** Heatmaps of sample-to-sample correlation for H2K27ac ChIP-seq replicates. Values indicate Pearson correlation. **E)** Scatter plot of normalized ATAC-seq signal at all distal open chromatin regions (> 3kb from TSS) in microglia after vehicle or IL-33 exposure. Color codes indicate significant changes (FDR < 0.05 & FC >2) in ATAC-seq signal (IL-33 enriched= red, vehicle enriched=black). **F)** Enriched *de novo* motifs in distal open chromatin regions (enhancers) that gain or lose H3K27ac ChIP-seq signal after treatment with IL-33 or vehicle, showing best matched TFs binding to those motifs (P_adj_ < 0.05). **G-H)** Browser tracks of ChIP-seq peaks for AP-1 family members ATF3, FOS and JunD in macrophages after LPS stimulation (from Fonseca et al., 2019^50^), shown above microglia ATAC-seq and H3K27ac ChIP-seq peaks 4 hours after IL-33 (this study). Highlighting *Marco* **(G)** and *Tlr2* **(H)**. Yellow shading denotes promoter regions, pink shading denotes distal gene regulatory elements (enhancers). **I)** qRT-PCR of *Fos* mRNA expression in cortical and thalamic microglia from indicated genotypes. Values were normalized to housekeeper (*Hmbs*) and control + vehicle condition. In Cx3cr1^creERT2^ control=Cx3cr1^creERT2+/-^. In P2ry12^creERT2^ control=*Il1rl1^fl/fl^.* Each dot represents a mouse. Two-way ANOVA followed by Tukey’s post hoc comparison (genotype and treatment). Data represented as mean ± SEM for bar graphs. \**p*<0.05, \*\*\**p*<0.001.

**Figure S5.**
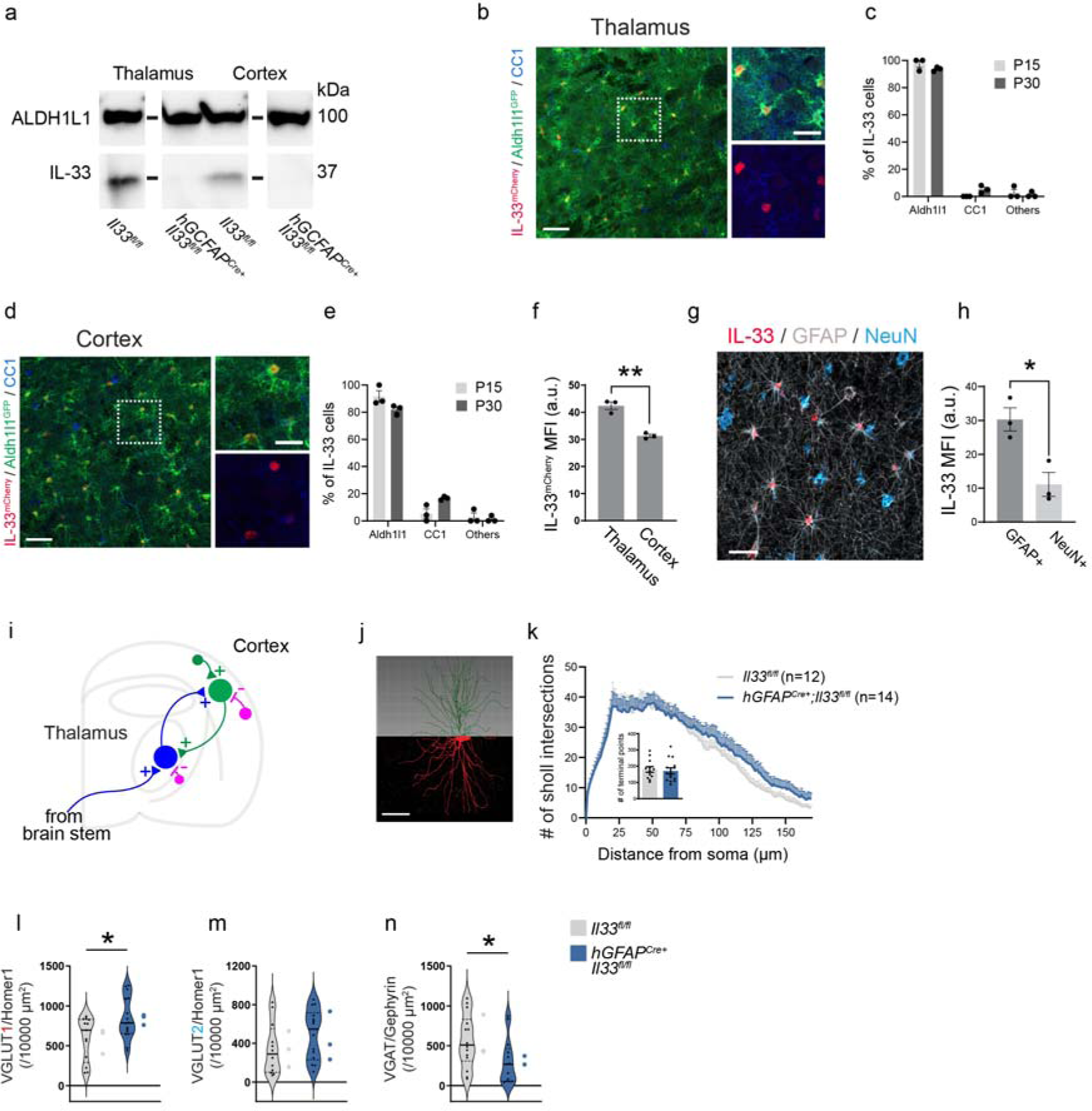
Defining the cellular sources of IL-33 in the developing thalamus and cortex, efficiency of IL-33 depletion using *hGFAPcre:Il33^fl/fl^*, and additional characterization of corticothalamic circuits after CNS-specific deletion of IL-33, related to Figure 4. **a)** Western blot from cortex and thalamus of *hGFAP^Cre+^:Il33^fl/fl^* animals and *Il33^fl/fl^* controls at P35. ALDH1L1 used as a loading control. Blot has been cropped to remove unrelated bands. **b-c)** Representative image and quantification of percent IL-33^mCherry^+ cells in the somatosensory thalamus of *Il33^mCherry^;Aldh1l1*^GFP^ mice stained with CC1 (oligodendrocytes) at P15 and P30. GFP expression marks astrocytes. Scale bar = 50 µm (left) and 20 µm (right). **e-e)** Representative image and quantification of percent IL-33^mCherry^+ cells in the somatosensory cortex of *Il33^mCherry^;Aldh1l1*^GFP^ mice stained with CC1 (oligodendrocytes) at P15 and P30. Scale bar = 50 µm (left) and 20 µm (right). **f)** Comparison of mean fluorescent intensity (MFI) of IL-33^mCherry^ in the thalamus and the cortex at P15. **g)** Representative image of IL-33 expression in the human cortical grey matter. Colocalized with GFAP (astrocytes) and NeuN (neurons). Scale bar = 20 µm. **h)** Mean fluorescence intensity of IL-33 expression in human cortical astrocytes and neurons. n= 3 male subjects aged 17, 50, and 51. **i)** Schematic of corticothalamic circuit. Green; VGLUT1+ excitatory neuron Blue; VGLUT2+ excitatory neuron. Purple; VGAT+ inhibitory neuron. **j)** Representative image and 3D reconstruction of biocytin filled neuron in somatosensory thalamus. **k)** Scholl analysis quantification of process branching and (inset) number of terminal branch points (two-way ANOVA followed by Sidak’s multiple comparison for Scholl analysis and two-tailed t-test for inset). Each dot represents a neuron. n=14 neurons from 7 *Il33^fl/lfl^* animals and n=12 neurons from 6 *hGFAPCre+:Il33^fl/fl^* animals. **l-n)** Quantification of synapses in somatosensory cortex in *Il33^fl/lfl^* and *hGFAPCre+:Il33^fl/fl^* animals, including excitatory intracortical **(l),** excitatory thalamocortical **(m),** and inhibitory **(n)** synapses. n=13 fields of view from 3 *Il33^fl/lfl^* and n= 16 fields of view from 3 *hGFAPCre+:Il33^fl/fl^* mice in D; n= 12 fields of view *Il33^fl/lfl^* mice and n= 14 fields of view from *hGFAPCre+:Il33^fl/fl^* mice in E; n= 13 fields of view from 3 *Il33^fl/lfl^* mice and n=14 fields of view from 3 *hGFAPCre+:Il33^fl/fl^* mice in F. Two-tailed unpaired t-test was used. Data represented as mean ± SEM for bar graphs and as median ± interquartile range for violin plots. Larger dots to the right of violin plots represent the average per individual mouse within that group. Mice from P26-P36 were used for k. Mice from P28-P30 were used for l-n. MFI; mean fluorescent intensity. \**p*<0.05, \*\**p*<0.01.

**Figure S6:**
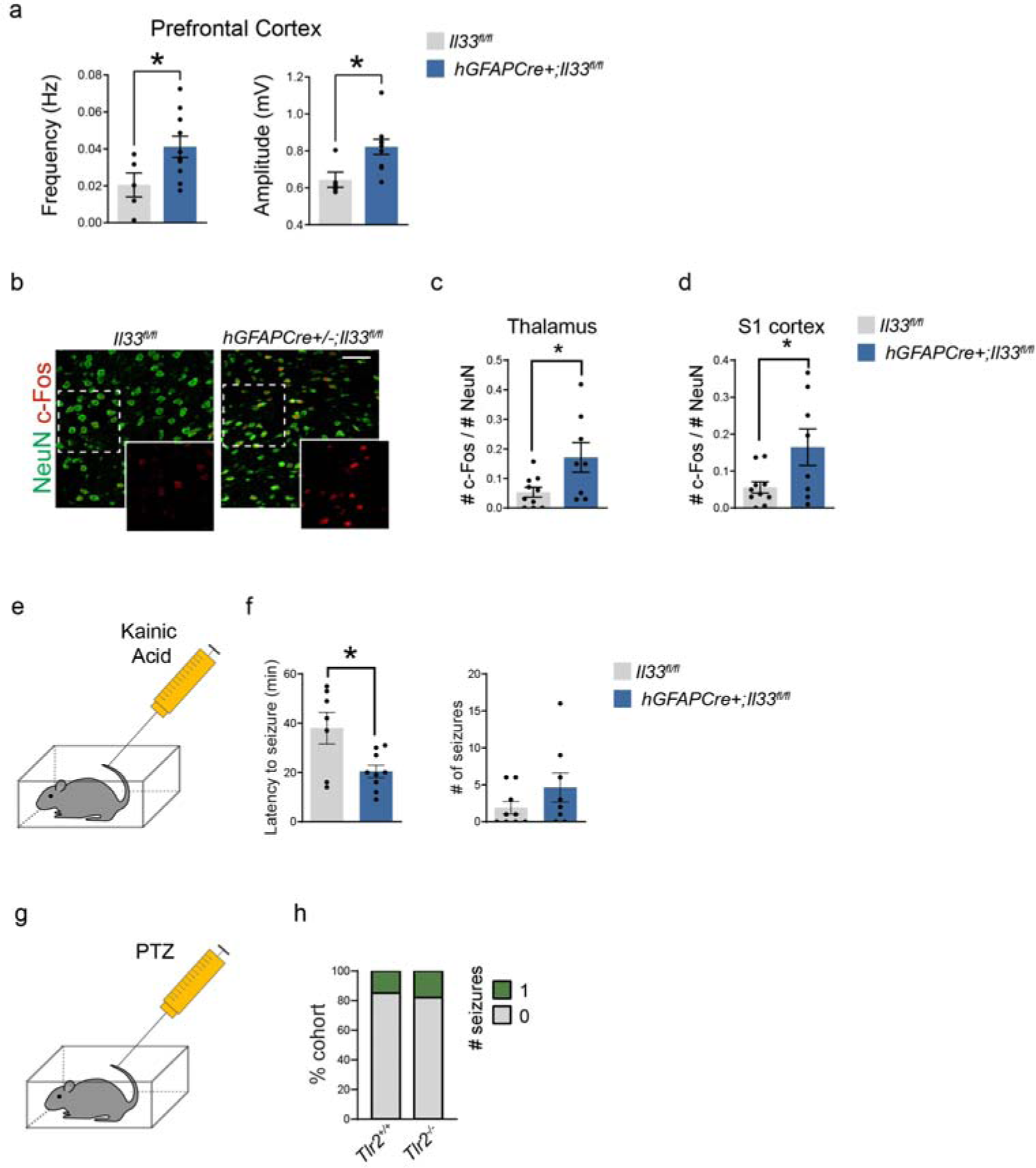
Additional characterization of seizure phenotypes after conditional deletion of IL-33, its receptor, and target genes, related to Figure 5. **a)** Quantification of total spike frequency (left) and average amplitude (right) of detected spike events from prefrontal cortex during one-hour recording session. n=10 *hGFAPCre^+^;Il33^fl/fl^* mice and 5 *Il33^fl/fl^* mice (two tailed t-test). Each dot represents a mouse. Mice were P35-P40. **b-d)** Representative images in thalamus **(b)** and quantification of c-Fos expression in the thalamus **(c)** and cortex **(d)** following PTZ administration (two-tailed t-test). Scale bar = 50 µm. Each dot represents a mouse. **e)** Schematic of kainic acid administration. **f)** Quantification of latency to first seizure onset (left) and incidence of seizures (right) for 3 hours following kainic acid administration (two-tailed t-test). Each dot represents a mouse. **g-i)** Quantification of percent of cohort with generalized tonic-clonic seizures in 1 hour following PTZ administration in wild-type vs. *Tlr2*^-/-^ animals. n=13 wild-type and 11 *Tlr2*^-/-^ animals from 2 independent experiments (Fisher’s exact test). Data represented as mean ± SEM for bar graph. Mice from P29-P35 were used for all experiment except A. \**p*<0.05.

**Table S1: ScRNA-seq data showing differentially expressed genes per cluster.** Data shows genes expressed in >5% of cells in that cluster, adjusted p-value < 0.001 and Log_2_ FC > 0.1/-0.1). See excel file.

**Table S2: Genes differentially expressed in microglia after IL-33 i.c.v. vs.** vehicle, in bulk RNA-seq (adjusted p-value < 0.01). **See excel file.**

**Table S3: ATAC-seq and H3K27ac ChIP-seq peaks differentially expressed in microglia after IL-33 i.c.v. vs. vehicle.** See excel file.

**Table S4:** mEPSC and mIPSC amplitude and kinetics, and intrinsic electrical membrane properties of neurons in somatosensory thalamus of IL-33 cKO (*hGFAPCre+/-^;^Il33^fl/fl^*) vs. littermate controls (*Il33^fl/fl^*) Data represented as mean±SEM. For mEPSC and intrinsic electrical membrane properties, *Il33^fl/fl^*: n=17 neurons from 3 mice, *hGFAPCre+/-^;^Il33^fl/fl^*: n= 18 neurons from 4 mice; For mIPSC, *Il33^fl/fl^*: n=16 neurons from 3 mice and *hGFAPCre+/-^;^Il33^fl/fl^*: n=17 neurons from 3 mice. Two-tailed unpaired t-test. ns: not significant.

| <u>mEPSC</u> | <u>Control</u> | <u>IL-33 cKO</u> | <u>p-value</u> |
| --- | --- | --- | --- |
| Amplitude (pA) | 12.43±0.40 | 11.76±0.40 | ns |
| Decay time constant (ms) | 2.77±0.13 | 2.88±0.22 | ns |
| Half-width (ms) | 1.52±0.09 | 1.53±0.12 | ns |
| <u>mIPSC</u> | <u>Control</u> | <u>IL-33 cKO</u> |  |
| Amplitude (pA) | 34.77±1.60 | 31.51±1.54 | ns |
| Decay time constant (ms) | 6.29±0.18 | 6.73±0.48 | ns |
| Half-width (ms) | 6.00±0.19 | 6.23±0.31 | ns |
| <u>Intrinsic electrical membrane properties</u> | <u>Control</u> | <u>IL-33 cKO</u> |  |
| Membrane capacitance (pF) | 125.48±6.23 | 131.70±7.04 | ns |
| Input resistance (mOhm) | 288.1±37.8 | 300±49.2 | ns |
| Resting membrane potential (mV) | -66.57±1.10 | -66.69±1.17 | ns |

## CONTACT FOR REAGENT AND RESOURCE SHARING

Anna Molofsky

## EXPERIMENTAL MODELS AND SUBJECT DETAILS

### Mice

All mouse strains were maintained in the University of California San Francisco specific pathogen–free animal facility, and all animal protocols were approved by and in accordance with the guidelines established by the Institutional Animal Care and Use Committee and Laboratory Animal Resource Center. Littermate controls were used for all experiments when feasible, and all mice were backcrossed >10 generations on a C57Bl/6 background unless otherwise indicated. The following mouse strains used are described in the table above and are as referenced in the text. For experiments using conditional alleles, Tamoxifen (Sigma, T5648) was diluted in corn oil (Sigma-Aldrich, C8267) at 37°C overnight and administered intragastrically at a concentration of 50 mg/kg three times every other day beginning at P1-P2 for Cx3cr1-creER and P2ry12- creER. 4-hydroxytamoxifen (Hello Bio, HB6040) was dissolved at 20 mg/mL in ethanol by shaking at 37°C for 15 min and was then aliquoted and stored at –20°C for up to several weeks. Before use, 4-OHT was redissolved in ethanol by shaking at 37°C for 15 min, and corn oil (Sigma-Aldrich, C8267) was added to give a final concentration of 2.5 mg/mL 4-OHT. The final 2.5 mg/mL 4-OHT solutions were always used on the day they were prepared and administered intraperitoneally at a concentration of 50 mg/kg.

## METHOD DETAILS

### Stereotaxic injections

All injections were performed with a Kopf stereotaxic apparatus (David Kopf, Tujunga, CA) with a microdispensing pump (World Precision Instruments) holding a beveled glass needle with ∼50 μm outer diameter. For perinatal experiments, mice were anesthetized by hypothermia. r all postnatal (>P8) and adult injections, mice were anesthetized with 1.5% isoflurane at an oxygen flow rate of 1L/min, headfixed with a stereotaxic frame, and treated with ophthalmic eye ointment. Fur was shaved and the incision site was sterilized with 70% ethanol and Betadine prior to surgical procedures. A hole was drilled in the skull. After injection, glass pipette was left in place for several minutes to allow diffusion. Pipette was slowly removed, and scalp re-apposed with tissue glue. Body temperature was maintained throughout surgery using a heating pad. Post-surgery Buprenorphine (Henry Schein Animal Health) was administered as needed by intraperitoneal injection at a concentration of 0.1 mg/kg. Further details for each experiment are below.

### FosTRAP2 labeling of Fos-expressing microglia

Homozygous Fos-TRAP2 mice (*Fos2A-CreER*) were crossed to R26R-lsl-TdTomato (*Ai14*) reporter mice. Progeny heterozygous for both alleles were administered 40 ng of recombinant IL-33 or vehicle (PBS) intracerebroventricularly as described separately. At one and four hours after i.c.v. injection, mice were injected intraperitoneally with freshly prepared 4-hydroxytamoxifen (4-OHT) a more rapidly bioavailable form of tamoxifen, at a concentration of 50 mg/kg.

Mice were sacrificed 24 hours after IL-33 i.c.v. injection. In a field of view, individual microglia IBA1 immunostaining was used to mask individual microglia, and Fos-CreER-Tdt positive and negative microglia were examined for TLR2 mean fluorescence intensity (as described separately) and presence or absence of MARCO staining.

### IL-33 and blocking antibody delivery

For bulk RNA, ATAC and H3K27ac ChIP sequencing, and quantitative RT-PCR, 500 ng of IL-33 or PBS (1 ) was slowly injected (3-5 nl/sec) into right lateral ventricle (ML = 1.25 mm, AP = -0.6 Z = -1.85 mm) of P30 mice. For all other experiments including single cell RNA-seq, *in vivo* microglia engulfment assay, *in vivo* microglia protein expression quantification, synapse counting, and FosTRAP2, either 40 ng of IL-33 or PBS (1 μl was slowly injected (3-5 nl/sec) into right lateral ventricle (ML = 1.1 mm, AP = 3.5 mm from lambda, Z = -1.8 mm) of P14- P16 mice. For MARCO and TLR2 antibody blockade either 0.8 µg of MARCO antibody (Bio-Rad) ^34, 66, 67^ or 1.6 µg of TLR2 antibody (Invivogen) or the same amount of IgG negative control antibody (Bio-Rad, MCA6004GA for MARCO and Invivogen, bgal- mab10-1 for TLR2) was administered in the same needle with IL-33.

### Quantification of mean fluorescence intensity of MARCO and TLR2

For *in vivo* microglia protein expression assay, mice were sacrificed 16-18 hours after IL-33 i.c.v. injection. For quantification of MARCO and TLR2 intensity, 4 um-thickness z-stack image was obtained using an LSM 800 confocal microscope (Zeiss) and maximum intensity projection image was created. Mean fluorescent intensity was quantified in ImageJ by first thresholding the Iba1 channel to make mask for microglia soma and process, then measuring the intensity of MARCO and TLR2 channel in masked area in grey scale and averaging those values in a field of view. Default thresholding was used for Iba1 masking.

### Fluorescence activated cell sorting (FACS) of microglia

Four hours after i.c.v. injection of IL-33 or PBS, P30 mice were anesthetized with isoflurane and perfused with PBS. As described ^68^, in brief, the brain was isolated and placed in ice-cold HBSS-Ca/Mg free supplemented with Hepes and glucose. The cortex was dissected and homogenized into a single cell suspension using a tissue homogenizer (5 cm^3^, VWR) and filtered m strainer (Falcon). Cells were pelleted at 300 xg for 10 min at 4°C and supernatant μ s discarded. A 22% Percoll gradient was run on the pellet to deplete wamyelin at 900 xg, no brake at 4°C and the pellet was afterwards incubated with CD16/CD32 (eBioscience), CD11b-PE (eBioscience) or CD11b-APC (BioLegend), CD45-FITC (eBioscience) or CD45-APC (BioLegend) and Ly-6C-APC or Ly-6C APC/Cy7 (Biolegend) antibodies in HBSS-Ca/Mg free supplemented with Hepes, glucose and EDTA (iMed-) on ice for 30 minutes. Cells were pelleted at 300 xg for 10 min at 4°C, resuspended in iMed- and incubated with DAPI just before FACS. A purified microglia population defined as CD11b^high^CD45^low^Ly-6C^neg^ was collected by FACS on a BD Aria3 (BD Biosciences). For scRNA sequencing a CD45+ population was collected as shown in supplementary figure 1A and processed further as described in 10x Genomics manual. For bulk RNA sequencing and qPCR, microglia were lysed with RLT+ (Qiagen) and stored at -80°C. For ATAC and H3K27ac ChIP sequencing microglia were processed as described below.

### Single-cell RNA sequencing of CD45+ cells

After FACS, approximately 10,000 CD45+ cells were loaded into each well of a 10x Genomics Chromium Chip G (v3.1) and dual-index libraries were prepared as described in the 10x Genomics manual. Library quality was assessed by Agilent High Sensitivity DNA kit on a Bioanalyzer (Agilent) and libraries were pooled and sequenced on Illumina NovaSeq SP100.

### Single-cell RNA sequencing analysis

Sequenced libraries were processed using the Cell Ranger 5.0 pipeline and aligned to the GRCm38 (mm10) mouse reference genome. Clustering and differential expression analysis were performed using Seurat version 4.0.1 ^69, 70^. Cells expressing fewer than 1300 unique genes and 2000 unique transcripts were excluded as likely debris, while cells expressing more than 5000 genes or 20,000 transcripts were excluded to remove cell doublets. Cells with higher than 5% mitochondrial transcripts were excluded to remove cells with membrane damage. Over 70% of the cells in each sample shown passed quality control thresholding for a total of >2000 cells per sample (*Cx3cr1^creERT2+/-^* + IL33 (Control + IL-33): 2205, *Cx3cr1^creERT2+/-^* + PBS (Control + PBS): 2730, *Cx3cr1^creERT2+/-^:Il1rl1^fl/fl^* + IL-33 (Il1rl1 cKO + IL33): 2707 healthy cells). An additional sample (*Cx3cr1^creERT2+/-^:Il1rl1^fl/fl^* + PBS) was excluded because less than half of the 1800 initially identified cells passed the quality control thresholds. Cells were identified as “female” or “male” based on their expression of the gene *Xist*; any cells expressing at least one count of *Xist* were labelled female, while all others were labelled male. The top seven transcripts correlated with sex (*Xist, Tsix, Ddx3y, Eif2s3y, Fkbp5, Ddit4, Uty*) were identified using the VariancePartition ^71^ package in R (1.20.0) and excluded from the PC, UMAP, and clustering calculations described below.

The top 6000 most variable genes, excluding the 7 sex-correlated genes above, were identified and their transcript counts normalized and scaled using the sctransform ^72^ function in Seurat, regressing out percent mitochondrial RNA and total counts per cell. 50 principal components were calculated from the scaled genes. The Harmony package (1.0) ^73^ was used to adjust the top 50 PCs to reduce technical variability between samples. These adjusted PCs were used for nearest neighbor, UMAP, and cluster identification. Cells were initially clustered with a resolution of 1, and two clusters (57 cells) with low expression of a canonical microglial gene (Cx3cr1) and nonzero expression of the myelin gene Mbp were excluded from downstream analysis as likely non-microglial contamination. The remaining cells were then passed through all of the normalization and clustering steps described in this paragraph again. A clustering resolution of 0.4 was used to generate 6 clusters.

Differential Expression (DE) analysis was done in Seurat using the MAST test ^74^ on the 6,000 most variable genes including only those genes expressed in at least 10% of the cells in a cluster. A p-value was calculated only for genes with a fold change of 5% or more.

Heatmaps were created with the DoHeatmap function in Seurat including the top 3 genes by log_2_ fold change per cluster, including only genes with an adjusted p-value lower than 10^-9^. 100 randomly selected cells are shown per cluster. Feature and Dimensional UMAP plots show 2,000 cells per sample. For the phagocytosis gene-specific heatmap, phagocytosis genes were identified using the GO term “Phagocytosis” (GO:0006909) and subset to include genes upregulated in cluster 1 by at least 7% (LFC>0.1), then ordered by descending log fold change in cluster 1. Only 100 cells are shown per cluster. Gene ontology (GO) analysis was performed using the Metascape webpage^75^(https://www.metascape.org) and only GO terms were used for Figure 1D.

### Bulk RNA sequencing of cortical microglia

RNA was isolated from 30000-60000 FACS-microglia per sample with the RNeasy® Plus Micro kit (Qiagen). Quality and concentration were determined with the Agilent RNA 6000 Pico kit on a Bioanalyzer (Agilent). All samples had an RNA Integrity Number (RIN) >7. cDNA and libraries were made using the Ovation® RNA-Seq System V2 kit (NuGen) and quality was assessed by Agilent High Sensitivity DNA kit on a Bioanalyzer (Agilent) and quantified by qPCR. Pooled libraries were RNA sequenced on an Illumina HiSeq 4000 paired-end for 125 cycles (PE125) yielding 50-70 million reads per sample.

### Bulk RNA sequencing analysis

Read quality was assessed with FastQC (http://www.bioinformatics.babraham.ac.uk /projects/fastqc) and aligned to the Mus Musculus genome (Ensembl GRCm38) using STAR aligner (version 2.6.0) ^76^, with the additional command --outFilterMultimapNmax 1, to only keep reads that map one time to the reference genome. Aligned reads were counted using HTSeq (version 0.9.0) ^77^ and counts were loaded into R (The R Foundation). DESeq2 package (version 1.24.0) ^78^ was used to normalize the raw counts and perform differential gene expression analysis.

### qPCR

RLT+ (Qiagen) lysed microglia were vortexed and frozen at -80° for storage. Samples were thawed and mRNA was isolated using RNeasy® Plus Micro kit (Qiagen). Purified mRNA was converted to cDNA using the High Capacity cDNA Reverse Transcription kit (Life Technologies). Primers for *Hmbs*, *Rps17*, *Marco*, *Tlr2*, *Il1rl1* and *Fos* were made using NCBI Primer Blast and ordered from IDT. A qPCR was run using Fast SYBR Green Master Mix (Thermo Fisher) on a 7900HT Fast Real-Time PCR System (Applied Biosystems). Data was analyzed using SDS software v2.4 (Applied Biosystems).

### Assay for transposase accessible chromatin (ATAC) sequencing of cortical microglia

Around 40000-70000 microglia were FACS isolated and collected into iMed-. Cells were pelleted at 300 xg, 4°C. Afterwards a previously published protocol ^44^ with modifications was used to perform ATAC sequencing. In brief, the pellet was gently resuspended in ice-cold 50 μl lysis buffer (10 mM Tris-HCl pH 7.4, 10 mM NaCl, 3 mM MgCl2, 0.1% IGEPAL C ^μ^ 630) and spun down at 500 xg for 10 min and 4°C. The supernatant was discarded, and cell pellet gently resuspended in 20 μ ltagment DNA buffer (Nextera, Illumina), 1 μtagment DNA enzyme (Nextera, Illumina), l nuclease free water) and incubated at 37 °C for 30 minutes. Samples were stored at -9 μ20 °C afterwards. The next day, tagmented DNA was purified using MinElute PCR purification kit (Qiagen) and size selected for 70 – 500 bp using AmpureXP beads (Beckman Coulter). Libraries were constructed and amplified using 1.25 μ Nextera index primers and NEBNext High-Fidelity 2x PCR Master Mix (New England BioLabs). A quantitative PCR was run to determine the optimal number of cycles. Libraries were afterwards twice size selected with AmpureXP beads (Beckman Coulter) for 150-400 bp fragments and paired-end sequenced for 100 cycles (PE100) on an Illumina HiSeq 4000 yielding 40-60 million reads per sample.

### ATAC sequencing analysis

FASTQ files were mapped to the mouse mm10 genome (UCSC). Bowtie2 with default parameters was used to map ATAC-seq experiments ^79^. HOMER was used to convert aligned reads into ‘tag directories” for further analysis ^80^. Peaks were called with HOMER for each tag directory with parameters -L 0 -C 0 -fdr 0.9 -minDist 200 -size 200. IDR was used to test for reproducibility between replicates, and only peaks with IDR < 0.05 were used for downstream analysis. The pooled tag directory from four replicates was used for track visualization using the UCSC genome browser^81^. To quantify chromatin accessibility, peak files were merged with HOMER’s mergePeaks and annotated with raw tag counts with HOMER’s annotatePeaks using parameters - noadj, -size given. DESeq2 ^78^ was used to identify chromatin accessibility with > 2 fold-change and adj. p-value < 0.05. Motif enrichment was performed using HOMER’s motif analysis (findMotifsGenome.pl) and de novo motifs was used. The background sequences were from the comparing condition as indicated in the figure legends.

### Chromatin immunoprecipitation-sequencing (ChIP-seq)

ChIP for H3K27ac was performed essentially as described previously ^82^. In brief, FACS purified cells were fixed with 1% formaldehyde for 10 min at room temperature. Next, formaldehyde was quenched with 2.625M glycine for 5 min at room temperature. Cells were collected by centrifugation at 1,500 x g for 10 min at 4°C. Cell pellets were then snap frozen and stored at -80C. For ChIP reactions, cell pellets were thawed on ice and lysed in 130 µl LB3 (10mMTris/HCl pH 7.5, 100mMNaCl, 1mMEDTA, 0.5mM EGTA, 0.1% deoxcycholate, 0.5% sarkosyl, 1 x protease inhibitor cocktail, and 1 mM sodium butyrate). Samples were sonicated in a 96 Place microTUBE Rack (Covaris cat#500282) using a Covaris E220 for 12 cycles with the following setting – time 60 s; duty 5.0; PIP, 140; cycles, 200; amplitude/velocity/ dwell 0.0. Samples were collected and 10% Triton X-100 was added to 1% final concentration. Sonicated samples were spun down at max speed, 4°C for 10 min. One percent of the sonicated lysate was saved as a ChIP input. The sonicated lysate was added to 20 µl Dynabeads Protein A with 1.5 ug anti-H3K27ac (Active Motif, #39685 Mouse Monoclonal) and incubated with slow rotation at 4°C overnight. The following day, beads were collected using a magnet and washed three times each with ice cold wash buffer I (20 mM Tris/HCl pH 7.5, 150 mM NaCl, 1% Triton X-100, 0.1% SDS, 2 mM EDTA, and 1 x protease inhibitor cocktail) and ice cold wash buffer III (10 mM Tris/HCl pH 7.5, 250 mM LiCl, 1% Triton X-100, 0.7% Deoxycholate, 1 mM EDTA, and 1 3 protease inhibitor cocktail). Beads were then washed twice with ice cold 10 mM Tris/HCl pH 7.5, 1 mM EDTA, 0.2% Tween-20. Sequencing libraries were prepared for ChIP products while bound to the Dynabeads Protein A initially suspended in 25 µl 10 mM Tris/HCl pH 8.0 and 0.05% Tween-20.

ChIP libraries were prepared while bound to Dynabeads using NEBNext Ultra II Library preparation kit with reaction volumes reduced by half, essentially as previously described (Heinz et al., 2018). Libraries were eluted and crosslinked reversed by adding to the 46.5 µl NEB reaction 20 ml water, 4 µl 10% SDS, 4.5 ml 5M NaCl, 3 ml 0.5 M EDTA, and 1 µl 20 mg/mL proteinase K, followed by incubation at 55C for 1 h and 65C overnight in a thermal cycler. Dynabeads were removed from the library using a magnet and libraries cleaned by adding 2 µl SpeedBeads 3 EDAC in 61 µl 20% PEG8000/1.5MNaCl, mixing well, then incubating at room temperature for 10 min. SpeedBeads were collected on a magnet and washed two times with 150 µl 80% ethanol for 30 s. Beads were collected and ethanol removed following each wash. After the second ethanol wash, beads were air-dried and DNA eluted in 25 µl 10 mM Tris/HCl pH 8.0 and 0.05% Tween-20. DNA was amplified by PCR for 14 cycles in a 50 ml reaction volume using NEBNext Ultra II PCR master mix and 0.5 mM each Solexa 1GA and Solexa 1GB primers. Libraries were cleaned up as described above using 2 µl SpeedBeads and 36.5 µl 20% PEG 8000/1.5 M NaCl and 2 µl SpeedBeads. After ethanol washing and drying, PCR amplified libraries were eluted from the SpeedBeads using 20 µl 10 mM Tris/HCl pH 8.0 and 0.05% Tween-20. Next, libraries were size selected 200-500 bp using gel extraction using 10% TBE acrylamide gels. Libraries were single-end sequenced using a HiSeq 4000.

### ChIP-sequencing Analysis

For preprocessing, Bowtie2 with default parameters was used to map ATAC-seq and ChIP-seq experiments^79^. HOMER was used to convert aligned reads into ‘‘tag directories’’ for further analysis^80^. To quantify H3K27ac signal, peak files were merged with HOMER’s mergePeaks and quantified with raw tag counts with HOMER’s annotatePeaks using parameters -noadj -size 1000 -pc 3 on ATAC-associated peaks. Peaks which contained at least 4 tags in at least 1 sample were used to identify differentially bounded peaks (FC > 2 and p-adj < 0.05) by DESeq2 ^78^. Peaks were categorized as distal peaks which are 3 kb away from known TSS. ChIP peak signals was normalized to the sequence depth. Motif Enrichment: To identify de novo motifs enriched in peak regions over random genomic background, HOMER’s motif analysis (findMotifsGenome.pl) was used. We performed a de novo motif analysis on H3K27ac signal within 300 bp of ATAC-associated peaks.

Data Visualization: The UCSC genome browser^81^ was used to visualize ChIP-seq and ATAC-seq data. The UCSC genome browser session containing the processed ATAC- and ChIP-seq data.

### Mouse immunohistochemistry

Mice were perfused transcardially with ∼10 mL of ice-cold 1X PBS followed by ∼10 mL of 4 % paraformaldehyde (PFA) diluted in PBS. Brains were post-fixed in 4 % PFA for a minimum of 3 hours and then transferred to a 30 % sucrose solution until brains fully submerged. Brains were then embedded in OCT compounds, frozen, and sliced in 14 µm thick coronal sections on a CryoStar NX70 Cryostat (ThermoFisher, MA). Brain sections were incubated in a blocking solution consisting of 5 % normal goat serum (ThermoFisher, MA) and 0.4 % Triton X-100 (Sigma-Aldrich) diluted in 1X PBS for 1 hour. Primary staining was done in the same blocking solution overnight at 4 °C. Secondary antibodies were diluted in the same blocking solution and tissue was incubated for 45 minutes at room temperature. Brain sections were mounted on coverslips with DAPI Fluoromount-G (SouthernBiotech, 0100-20) for all other experiments. Histology slides were imaged on an LSM780 or LSM800 or LSM880 confocal microscope with AiryScan (Zeiss, Germany) on Superresolution mode using a 63x objective for synapse quantification and microglia engulfment assay and an LSM 700 or LSM 800 confocal microscope (Zeiss, Germany) using 20x objectives for all other imaging. The following goat secondary antibodies and their dilutions were used corresponding to the host of primary antibodies; Alexa Flour 405 (1:500; Abcam), Alexa Fluor 488, Alexa Fluor 555, Alexa Fluor 647 (1:500; ThermoFisher).

### Human brain immunohistochemistry

Human brain tissue was collected during autopsy with postmortem interval < 48 hours. Tissue was collected with previous patient consent in strict observance of legal and institutional ethical regulations in accordance with the University of California San Francisco Committee on Human Research. Brains were cut into ∼1.5-cm coronal or sagittal blocks, fixed in 4% paraformaldehyde for 2 d, cryoprotected in a 30% sucrose solution, and embedded in optimal cutting temperature (OCT) compound (Tissue-Tek). Samples contained no evidence of brain disease as assessed by a neuropathologist (E.J. Huang). We collected 14-μm cryosections on Superfrost slides (VWR) using a Leica CM3050S cryostat. Secti μ s were allowed to equilibrate to room temperature for 3 hours. Antigen retrieval was conducted at 95°C in 10 mM sodium citrate buffer, pH 6. Following antigen retrieval, slides were washed with TNT buffer (0.05% Triton-X100 in PBS) for 10 minutes, placed in 0.5 % H_2_O_2_ in PBS for 30 minutes and then blocked with TNB solution (0.1M Tris-HCl, pH 7.5, 0.15M NaCl, 0.5% blocking reagent from PerkinElmer, NEL701A001KT) for 1 hour. Slides were incubated in primary antibodies overnight at 4°C and in secondary antibodies for 1 hour at room temperature. All antibodies were diluted in TNB solution. Sections were then incubated for 30 minutes in streptavidin–horseradish peroxidase, which was diluted (1:1250) with TNB. Tyramide signal amplification (PerkinElmer, NEL744001KT) was used for some antigens. Sections were incubated in tyramide-conjugated Cy3 (1:100) for 4 minutes. The following secondary antibodies and their dilutions were used; donkey anti-goat biotinylated antibody (1:500; Jackson ImmunoResearch, 705-065-147), donkey anti-chicken DyLight 405-conjugated antibody (1:500; Jackson ImmunoResearch, 703- 475-155), and donkey anti-rat Alexa Fluor 647-conjugated antibody (1:500; Jackson ImmunoResearch, 712-605-153). MaxEntropy thresholding was used for masking.

### Western blotting

Tissues were flash frozen on dry ice, then sonicated for 20 seconds in lysis buffer (50 mM tris-HCl, 1 mM EDTA, 1% Tx-100, 150 mM NaCl). The sample was centrifuged for 10 minutes at 15,000 rpm at 4L and the pellet was discarded. Samples were run on a denaturing gel and transferred to PVDF membrane, blocked with 5% milk in TBST for 1 hour at room temperature, incubated in primary antibody overnight at 4°C and secondary at room temperature for one hour, and developed with ECL plus. The following secondary antibodies and their dilutions were used; rabbit anti-goat HRP-linked secondary (1:1000, Bio-Rad, 1721034) and goat anti-rabbit HRP-linked secondary (1:2000, Cell Signaling Technology, 7074S).

### Slice preparation

Slices were prepared as previously described^12^. Briefly, we prepared 250 µm (for patch-clamp electrophysiology) or 400 µm (for *ex vivo* microglia engulfment assay)-thick horizontal slices including thalamus in ice-cold sucrose cutting solution containing 234 mM sucrose, 2.5 mM KCl,1.25 mM NaH2PO4, 10 mM MgSO4, 0.5 mM CaCl2, 26 mM NaHCO3, and 10 mM glucose, equilibrated with 95% O2 and 5% CO2, pH 7.4, using a Leica VT1200 vibrating microtome (Leica Microsystems) from 4 months old mice. We incubated the thalamic slices, initially at 32-34L for an hour and then at room temperature (for patch-clamp electrophysiology) or 35-36 engulfment assay), in artificial cerebrospinal fluid (aCSF) containing L (for microglia mM NaCl, 2.5mM KCl,1.25 mM NaH2PO4, 1 mM MgCl2, 2 mM CaCl2, 26 mM NaHCO3, and 10 mM glucose, equilibrated with 95% O2 and 5% CO2, pH 7.4.

### Whole cell patch-clamp recording

Recordings were performed as described^12^. Briefly, recording electrodes made of borosilicate glass had a resistance 3-5 MOhm when filled with intracellular solution containing 115 mM potassium gluconate, 11 mM KCl, 1 mM MgCl_2_, 1 mM CaCl_2_, 10 mM HEPES, and 11 mM EGTA, 2 mM K_2_ATP, 0.1% biocytin, pH adjusted to 7.35 with KOH (286 mOsm) for miniature excitatory post-synaptic currents (mEPSCs) recording or 129 mM CsCl, 5 mM QX-314Cl, 2 mM MgCl2, 10 mM HEPES, and 10 mM EGTA, 4mM MgATP, 0.1% biocytin, pH adjusted to 7.38 with CsOH (288 mOsm) for inhibitory post-synaptic currents (mIPSCs) recording. Series resistance was monitored in all recordings, and the recordings were excluded from analysis if the series resistance was > 25 MOhm or varied by more than 20%. Recordings were obtained using an MultiClamp 700B (Molecular Devices, CA), digitized (Digidata 1550B; Molecular Devices), and acquired at 20 kHz using the pClamp 10 software (Molecular Devices). Recordings were performed in voltage-clamp mode at a holding potential of -70 mV and obtained from visually identified neurons in somatosensory thalamus for 10 minutes. In the presence of 0.5 µm tetrodotoxin, 50 µm picrotoxin (Sigma-Aldrich, P1675) or 50 µm D-(-)-2-Amino-5-phosphonopentanoic acid (5AP; Hello Bio, HB0225) and 20 µm 6,7-Dinitroquinoxaline-2,3(1H,4H)-dione (DNQX, Sigma-Aldrich, D0540) were used to isolate mEPSCs or mIPSCs, respectively. The recordings were analyzed using ClampFit (Molecular Devices) and MiniAnalysis (Synaptosoft, NJ).

### Microglial engulfment assays

For quantification of *in vivo* microglial synapse engulfment, 16-18 hours after i.c.v. injection of IL-33 or PBS and MARCO antibody, TLR2 antibody or IgG negative control antibody. The brains were post-fixed in 4% PFA for a minimum of 6 hours and transferred to a 30% sucrose solution until brains fully submerged. The brains were cut in 30 um thick horizontal sections using HM 440E freezing microtome (GMI Instruments).

Quantification of engulfment was performed as previously described^13^ Briefly, Z-stacks encompassing entire microglia were collected with a Zeiss LSM880 confocal microscope with AiryScan (Zeiss, Germany) on Superresolution mode (∼150nm resolution) using a 63x objective, with NA 1.4. Laser power and gain were consistent across all experiments. Images were analyzed using Imaris software (Bitplane) by creating a 3D surface rendering of microglia, thresholded in pilot experiments to ensure that microglial processes were accurately reconstructed, and maintained consistent thereafter. This rendering of microglia was used to mask the CD68 channel to create a 3D surface rendering of phagolysosomes. This rendering was used to mask the VGLUT1 or Homer1 signal, and the “Spots” function was subsequently used to quantify the number of VGLUT1 puncta or Homer1 puncta entirely within the microglial surface or the surface of phagolysosome in individual microglia. Analysis was automated and experimenter was blinded to genotype and experimental condition throughout. In some cases, data was normalized to the indicated control conditions to better allow comparisons between different experimental batches.

### Quantification of c-Fos+ neurons

c-Fos+ neuron was quantified in ImageJ by first thresholding the NeuN channel to make mask for an individual neuron, thresholding the c-Fos channel, and analyzing the c-Fos+ neuron based on the NeuN mask image. Default thresholding was used for NeuN masking and MaxEntropy thresholding was used for c-Fos+ identification.

### IL-33^mCherry^ quantification

In most cases, positive and negative cells were quantified. For intensity measurements 7.5 um-thickness z-stack image was obtained using an LSM 800 confocal microscope (Zeiss) and maximum intensity projection image was created Mean fluorescent intensity was quantified in ImageJ by first thresholding the IL-33 channel to make mask for an individual cell, then measuring the intensity of IL-33 channel of an individual cell in grey scale and averaging those values in a field of view.

### Seizure behavior Assays

Mice between P30-P40 were used for seizure behavioral assays.

Pentylenetetrazol (PTZ; Sigma-Aldrich, MO) and Kainic acid (KA; Tocris, United Kingdom) were dissolved in normal saline and freshly prepared. 50 mg/kg of PTZ or 16 mg/kg of KA was used. Each animal was placed in the center of a transparent cage immediately after IP injection of PTZ or KA, and behavior was video recorded for 1 hour or 3 hours, respectively. Video clips were analyzed to measure the latency to develop seizures, defined as generalized tonic-clonic seizure (GTC) with loss of balance or wild jumping, the number and duration of seizures, scored on a Racine scale as previously described^83^. More than 10 seconds between two GTCs was considered as two separate events. After KA injection, 2/9 and 1/10 mice from control and IL33 cKO, respectively, which did not have a seizure, were excluded from the latency to seizure onset analysis, and 1/9 and 2/10 mice from control and IL33 cKO, respectively, which had continuous prolonged seizure and died, were excluded from duration and the number of seizures analysis. Mice were sacrificed within 3 hours after PTZ injection for IHC experiment. All experiments were conducted in the same conditions between 10AM and 4PM, and experimenter was blinded to genotype throughout data collection and analysis.

### Implantation of electrocorticogram (ECoG) devices

Procedures were performed as previously described ^84^. Custom devices containing multiple screws were used to acquire electrocorticogram (ECoG) signals *in vivo* (Mill-Max Manufacturing Corp, NY). Animals were anesthetized with isoflurane (2-5%) and secured with ear bars in a stereotaxic frame while resting on top of a small heating pad to maintain body temperature. Small bur holes were drilled using a hand-held drill (Dremel, WI), and were located above the prefrontal cortex (AP: +0.5 mm from Bregma) and somatosensory cortex (AP: -0.5 mm from Bregma, ML: +2.5 mm). The reference screw was placed above the cerebellum (AP: -1.0 mm from Lambda; ML: +1.0 mm). To implant ECoG devices, screws were placed into the bur holes and secured with dental cement. Topical lidocaine and antiseptic ointment were applied to the skin surrounding the implant. Animals were monitored for 1 week as they recovered, prior to beginning recordings.

### *In vivo* ECoG acquisition

Recordings were performed as previously described^84, 85^. ECoG signals were recorded at 24.41 kHz sampling rate using RZ5 and Synapse software (Tucker Davis Technologies). A video camera mounted on a flexible arm was used to continuously monitor the animals. Each recording trial consists of 60 minutes of baseline recording followed by 60 minutes of recording after a single-dose 50 mg/kg PTZ intraperitoneal injection. ECoG signals from prefrontal cortex and somatosensory cortex were referenced to the ECoG screw electrode placed over the cerebellum. Analysis was performed using ClampFit (Molecular Devices, CA) for spike and spike-wave discharge (SWD) analysis.

For spike analysis, data were bandpass filtered at 1 and 100 Hz. A spike was defined as a signal which was greater than 0.4 mV based on visual inspection of all recordings (mean standard deviation from mean spike amplitude (1 ± 3.6 µV) was 0.068 ± 0.015 mV).

For SWD analysis, data were bandpass filtered at 1 and 60 Hz. SWDs were defined as at least 5 connected rhythmic 4–6 Hz spike-wave complexes (typically spanning at least 0.5 seconds) with amplitudes at least two-fold higher than background. All data was collected and analyzed blinded to genotype and condition.

### Synapse quantification

Mice between P14-P17 were used for synapse quantification after i.c.v. IL-33 injection. Mice between P28-P35 were used for synapse quantification of *hGFAPCre*;*Il33^floxed^* and *Cx3cr1-creER;Il1rl1^floxed^* and their control experiments. Quantification was performed as previously described^43^. Briefly, synapse colocalization for pre-and post-synaptic markers was quantified by determining colocalization of Homer1 and VGLUT1 or VGLUT2; Gephyrin and VGAT in optical sections of somatosensory thalamus and layer 4 sensory cortex. Images were collected using standardized imaging parameters throughout, and colocalization was analyzed using PunctaAnalyzer, an ImageJ plug-in developed by the Eroglu lab. Analysis parameters were constant throughout and blinded to genotype and condition. Image quality was checked by repeating analyses after 90° rotation of one channel to verify that co-localization was not due to random chance.

### Dendritic branch quantification

For quantification of dendrite branch, biocytin-labeled 250 µm thickness acute brain slices were fixed in 4 % PFA overnight after whole-cell patch clamp recording, and then washed with PBS 3 times. Slices were incubated in Alexa Fluor 555-conjugated streptavidin (1:1000; ThermoFisher, S32355) at 4 L overnight in the dark. Z-stacks encompassing entire neuron were collected; Images were analyzed using Imaris software (Bitplane) by creating a filament rendering of neuron, manually corrected by blinded experimenter to ensure neuronal processes were accurately reconstructed.

### Statistical Analysis

For statistical analysis, Graphpad Prism 8, 9 and R was used. Comparisons of two groups were performed with the two tailed t-test, the nonparametric Mann–Whitney test, or the Fisher’s exact test, as needed. The difference between multiple groups was tested by one-way ANOVA followed by Tukey’s test. Two-way ANOVA followed by Tukey’s multiple comparison, Newman-Keuls test or Sidak’s multiple comparison was used as needed when more than two comparisons were made. The level of significance was set at *p* < 0.05. RNA-sequencing and ATAC-seq data were analyzed in R as described in the methods section above.

### Data and Materials Availability

RNA, ATAC, H3K27ac ChIP and scRNA sequencing data of microglia post i.c.v injection of IL-33 or PBS are available through GEO (number pending). Marco KO mice were kindly provided by Dr. James Luyendyk under MTA AAGR2021-00156.

## RESOURCE TABLE

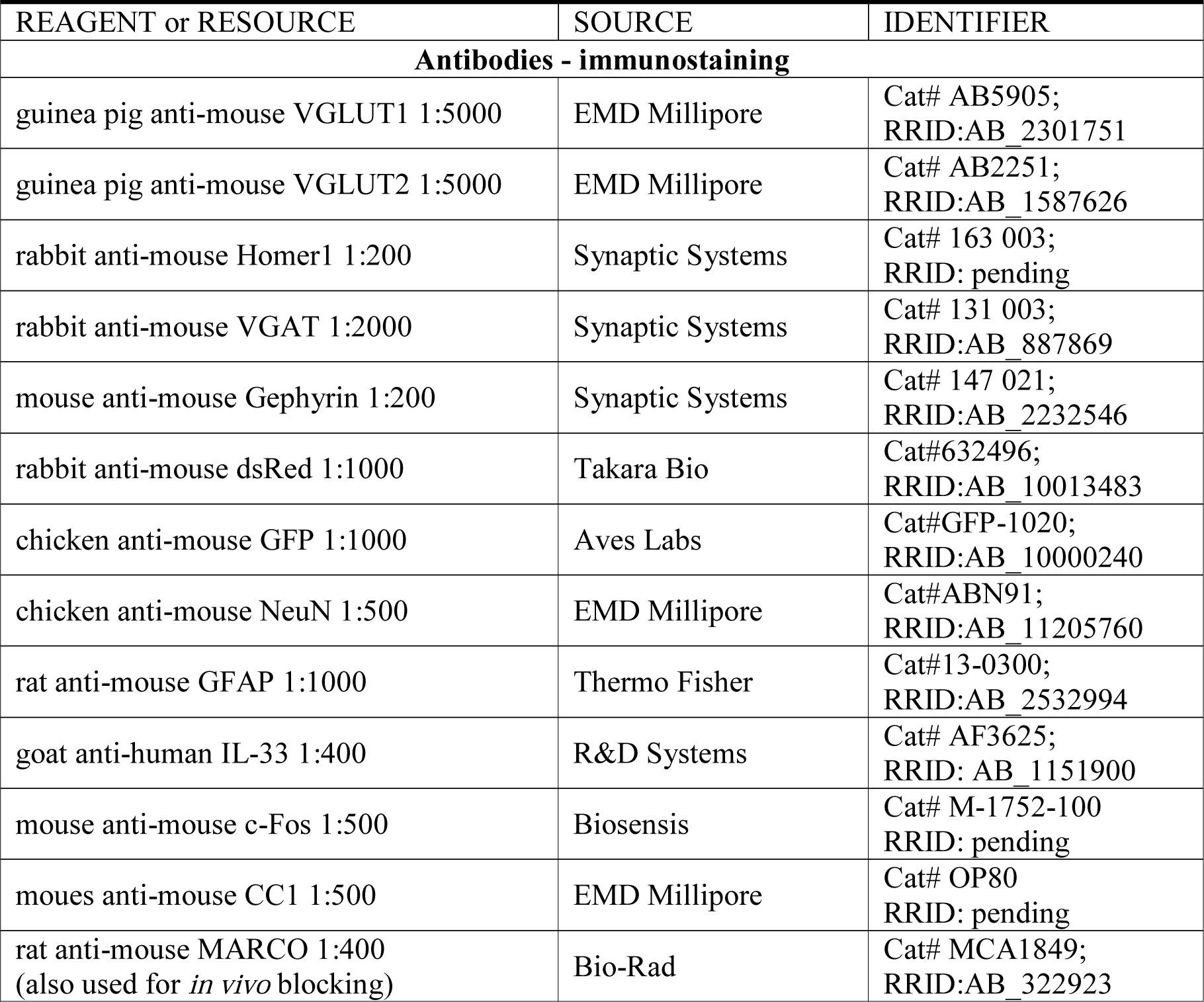
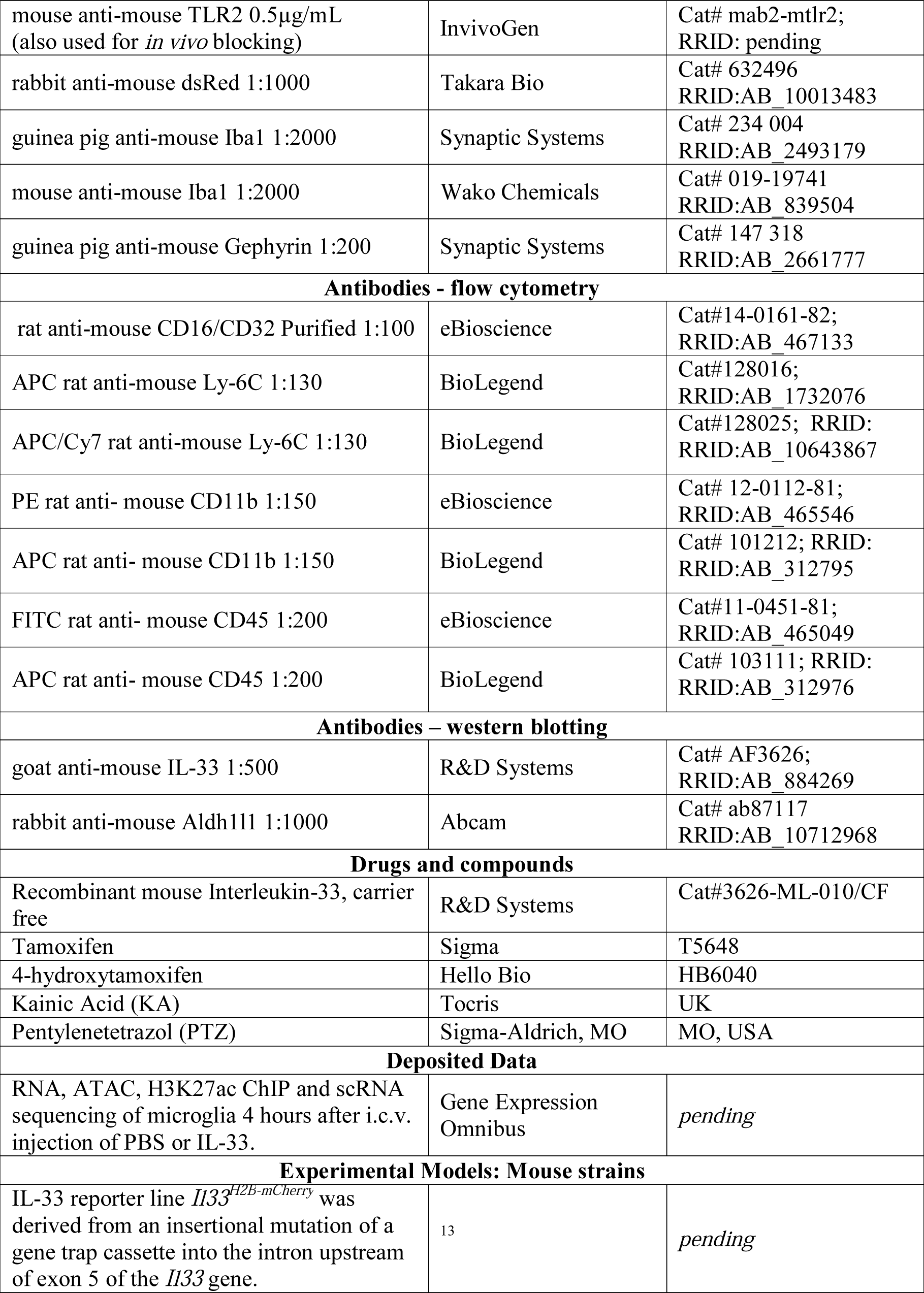
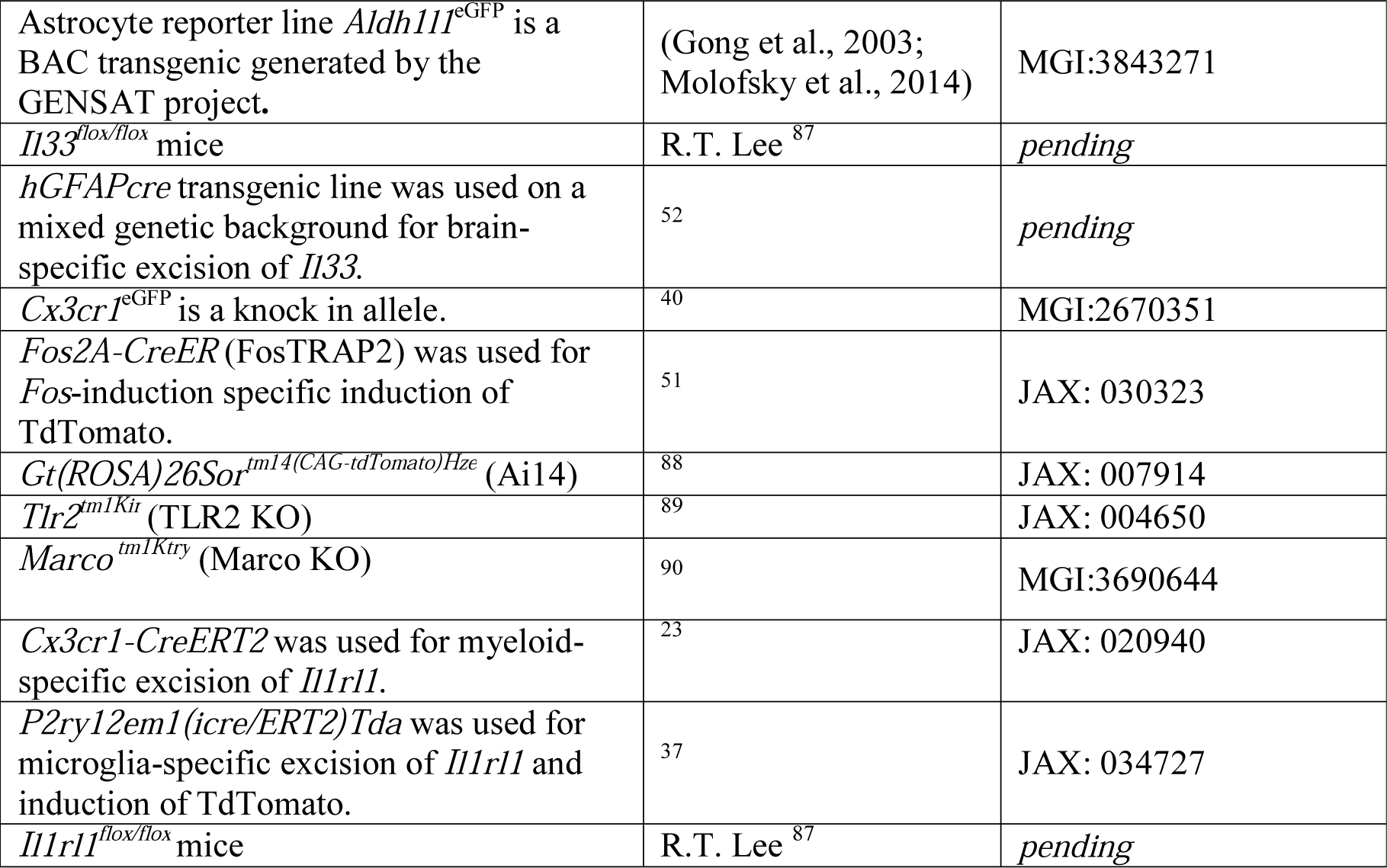

## Notes

### Competing Interest Statement

The authors have declared no competing interest.

